# Divergent clonal differentiation trajectories of T cell exhaustion

**DOI:** 10.1101/2021.12.16.472900

**Authors:** Bence Daniel, Kathryn E. Yost, Katalin Sandor, Yu Xia, Yanyan Qi, Kamir J. Hiam-Galvez, Stefanie L. Meier, Julia A. Belk, Josephine R. Giles, E. John Wherry, Howard Y. Chang, Takeshi Egawa, Ansuman T. Satpathy

## Abstract

T cells activated by chronic antigen exposure in the setting of viral infections or cancer can adopt an exhausted T cell (Tex) state, characterized by reduced effector function and proliferative capacity, and the upregulation of inhibitory receptors. However, whether all antigen-specific T cell clones follow the same molecular and cellular Tex differentiation trajectory remains unclear. Here, we generate a single-cell multi-omic atlas of T cell exhaustion that redefines the phenotypic diversity and molecular regulation of Tex phenotypes. Longitudinal analysis during chronic viral infection identifies an early effector phenotype that is epigenetically primed for Tex differentiation and two late-stage Tex cell states with either a terminal exhaustion or a killer cell lectin-like receptor (KLR)-expressing cytotoxic gene signature. We define clonal trajectories of antigen-specific T cells using paired single-cell RNA and T cell receptor sequencing and reveal distinct differentiation trajectories resulting in terminal Tex-biased, KLR Tex-biased, or divergent clones that differentiate into both phenotypes. Comparison of Tex phenotypes among shared T cell clones that traffic to multiple organs reveals that clonal differentiation trajectories are maintained across tissues. Finally, we show that differences in clonal differentiation trajectory are driven by TCR signal strength, whereby high-affinity T cell clones preferentially adopt a terminal Tex fate, while low-affinity clones adopt an effector-like KLR Tex fate that is detectable long-term but depleted in high antigen settings. These findings reveal clonal heterogeneity in the T cell response to chronic antigen and genomic programs that underlie Tex fates and persistence.

**Highlights:**

- A single-cell atlas of T cell exhaustion identifies novel early effector and KLR Tex states.
- Clonal T cell analysis defines divergent differentiation trajectories during chronic viral infection leading to terminal and KLR Tex fates.
- The heterogeneity of the Tex pool arises from three primary differentiation patterns and are differentially persistent in the setting of high antigen.
- Clonal Tex differentiation patterns are conserved across organ sites and driven by TCR signal strength.

## Introduction

Chronic antigen exposure during chronic viral infections and cancer leads to impaired CD8^+^ T cell responses, termed T cell exhaustion [1, 2]. Exhausted T cells (Tex) are characterized by reduced effector function, diminished proliferative capacity, and high expression of inhibitory receptors, including PD-1, LAG-3, and TIM3. However, Tex are able to maintain some effector functions and persist long-term, suggesting that T cell exhaustion may represent a mechanism to control pathogen burden while maintaining immune homeostasis [3, 4]. Recent studies have identified heterogeneity in Tex phenotypes, which are characterized by distinct surface receptors, functionality, proliferative capacity, and tissue localization during chronic viral infections and cancer [5–12]. Some of these studies have proposed a linear differentiation model, whereby progenitor Tex (Tex^prog^; marked by expression of TCF1 and CXCR5) self-renew and maintain downstream Tex subsets, including CX3CR1^+^PD-1^+^ intermediate Tex (Tex^int^) with proliferative, cytolytic and memory potential, and PD-1^+^ TIM3^+^ terminal Tex (Tex^term^; marked by high expression of inhibitory receptors, and limited effector or proliferative potential) [5–9, 13, 14, 60]. These subpopulations exhibit distinct epigenetic states and rely on distinct transcription factors (TFs). TCF1 (encoded by *Tcf7*) and BACH2 are indispensable for the formation of the Tex^prog^ phenotype, while the high mobility group TF, TOX, orchestrates the establishment and maintenance of the molecular program of exhaustion in all Tex states, including Tex^term^, and is required for their survival [4, 7, 15–19]. Finally, these Tex populations are further distinguished by their ability to respond to immune checkpoint blockade (ICB); Tex^term^ possess a stable epigenetic program and cannot be efficiently reinvigorated by ICB, while Tex^prog^ can proliferate in response to ICB and may be important for the therapeutic responses [3, 6, 20]. Despite these advances, we lack a comprehensive view of Tex states, their clonal relationships, and the molecular programming underlying their differentiation, particularly in polyclonal T cell responses.

Here, we generate a comprehensive atlas of Tex differentiation using single-cell chromatin accessibility, transcriptome, and T cell receptor (TCR) sequencing of antigen-specific CD8^+^ T cells in the setting of chronic lymphocytic choriomeningitis virus (LCMV) infection. We discover previously unappreciated Tex subsets, including an early effector exhausted subset (Tex^eeff^) in the early phase of infection that initiates the molecular program of exhaustion, and a killer cell lectin-like receptor-expressing Tex subset (Tex^KLR^) as a late-stage phenotype concurrent with terminal Tex, which suggests a divergent developmental path during Tex differentiation. T cell clone tracing based on paired scRNA/TCR-seq nominates unexpected diversity in Tex differentiation trajectories; namely, chronic antigen-specific T cell clones can adopt Tex^term^-biased, Tex^KLR^-biased, or divergent fates, comprising both cell types. Multi-organ clonal analysis reveals that Tex clones traffic to multiple organ sites and that their differentiation trajectories are conserved across tissues; however, Tex^KLR^-biased clones are depleted in the liver, suggesting that Tex^term^ cells may be phenotypically adapted for high-antigen tissue microenvironments. Finally, we show that clone behaviors are programmed by TCR affinity to cognate antigen; high-affinity TCR clones are biased towards a Tex^term^ differentiation trajectory, while low-affinity TCR clones are biased towards a Tex^KLR^ trajectory. Overall, these results provide an in-depth view of the gene regulatory programs and clonal dynamics of Tex states during chronic infection and suggest that a polyclonal T cell response to chronic antigen may balance T cell states that perform effector and memory functions.

## Results

### A multi-omic single-cell atlas of CD8^+^ T cell differentiation during acute and chronic viral infection

To profile CD8^+^ T cell differentiation during T cell exhaustion, we used mouse models of acute (LCMV Armstrong – Arm) or chronic (LCMV Clone 13 – Cl13) viral infection. These two viral strains only differ by two amino acids, and the immunodominant epitopes are identical, enabling direct comparisons of antigen-specific T cell responses in both models [21, 22]. We generated paired single-cell RNA- and T cell receptor (TCR)-sequencing (scRNA/TCR-seq) and single-cell assay for transposase accessible chromatin with sequencing (scATAC-seq) data from LCMV glycoprotein 33-41 tetramer positive (gp33^+^) and tetramer negative (gp33^-^) splenic CD8^+^ T cells at two timepoints (Day 8 and Day 21 post-infection) for both infection models (**Figure 1A-C**). At Day 21 (D21) of Cl13 infection, we also generated scRNA/TCR-seq of gp33^+^ and gp33^-^ populations from two additional organs with known differences in viral antigen load (lung and liver; **Figure 1B and Figure S1A**) [23, 24]. Finally, we sorted D21 Cl13 splenic T cells using previously defined surface markers that identify Tex^prog^ (SLAMF6^+^), Tex^int^ (CX3CR1^+^), and Tex^term^ (PD-1^+^, SLAMF6^-^ and CX3CR1^-^) phenotypes and performed scRNA/TCR- and scATAC-seq (**Figure S1B**) [7–9, 13]. In total, we obtained 96,750 scRNA-seq profiles that passed quality control filters based on the detected gene count (>200 genes/cell), mitochondrial content (<5% mitochondrial RNA content/cell), and predicted doublets (**Figure 1D and S1C, Methods**). Of scRNA-seq profiles passing quality control filters, we detected TCR alpha and beta sequences in 88,696 T cells (91.7%) and 5,197 expanded T cell clones (clones >1 cell; **Figure 1D**). We obtained 62,731 scATAC-seq profiles that passed quality control filters based on the unique ATAC-seq fragment count (>1,000 fragments/cell), median read enrichment at transcription start sites (>4 TSS score), and predicted doublets (**Figure 1E, S1D and S1E, Methods**).

**Figure 1.**
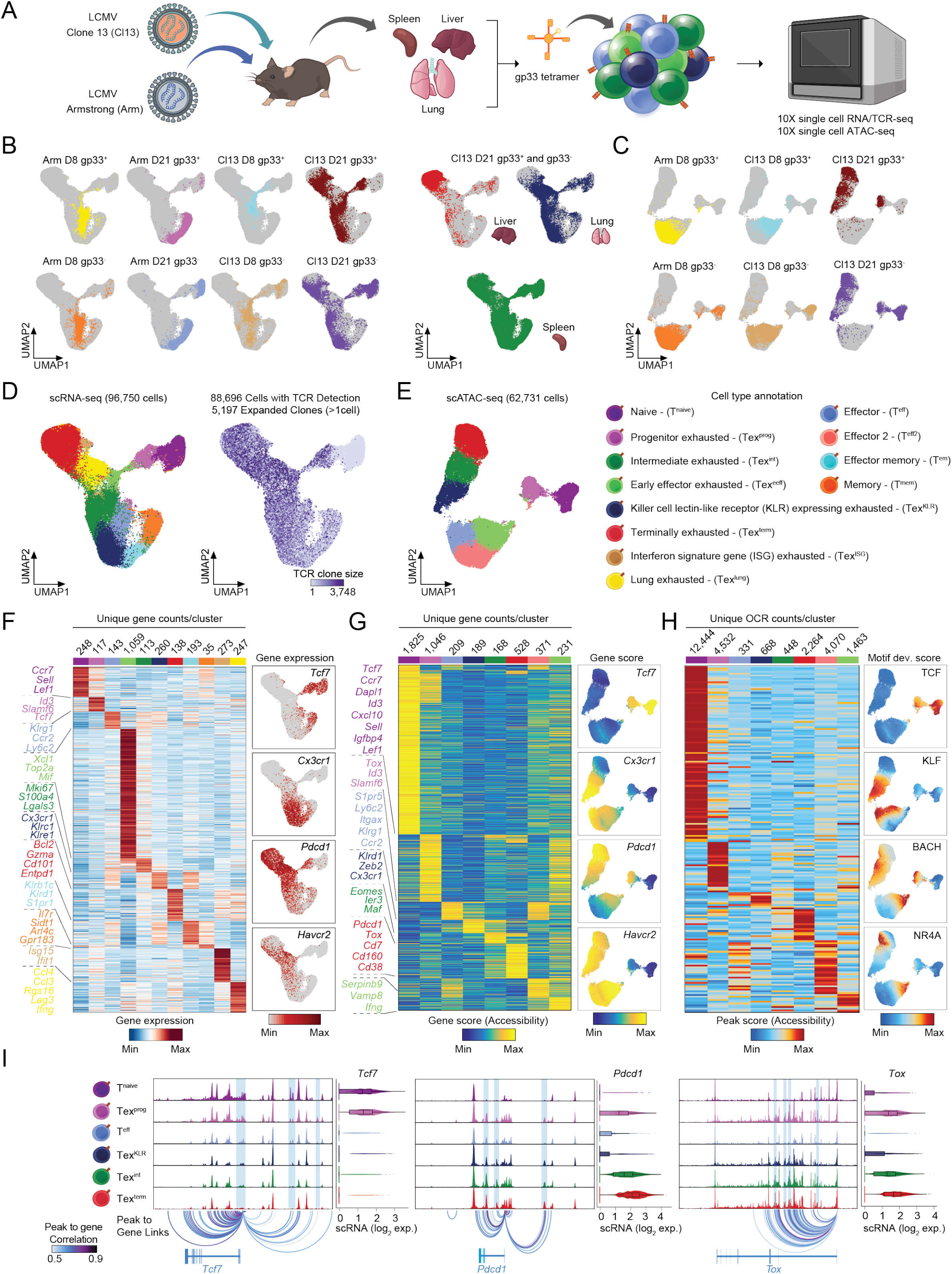
Single-cell genomic atlas of T cell exhaustion during LCMV infection. **(A)** Schematics on the mouse model used, indicating the two viral strains, the tetramer sort and the single cell technologies applied. **(B)** UMAPs of scRNA-seq profiles colored by the samples (gp33^+^ and gp33^−^ fractions) sorted from the spleen of Arm- or Cl13-infected animals on the indicated days (D8 and D21) (left). UMAPs of scRNA-seq profiles colored by the samples originating from the different organs of Cl13-infected animals at D21 (right). **(C)** UMAPs of scATAC-seq profiles colored by the samples (gp33^+^ and gp33^−^ fractions) sorted from the spleen of Arm- or Cl13-infected animals on the indicated days. **(D)** UMAP of all scRNA-seq profiles, colored by the annotated T cell subsets (left). UMAP of scTCR-seq results colored by the size of the expanded clones from which individual T cells originate (right). **(E)** UMAP of all scATAC-seq profiles colored by the annotated T cell subsets. **(F)** Heat map of subset specific marker genes determined by scRNA-seq. Feature plots of specific gene markers that characterize T cell subsets. **(G)** Heat map of Gene score values (accessibility) determined by scATAC-seq. Feature plots of specific Gene score values that mark main T cell subsets. **(H)** Heat map of Peak score values at the unique open chromatin regions (OCRs) of the T cell subsets determined by scATAC-seq. Feature plots show the motifs that are accessible in specific T cell subsets (chromVAR deviation scores are depicted). **(I)** Genome browser snapshots on the indicated gene loci, showing the chromatin states of the different T cell subsets. Violin plots show the associated expression level of the indicated genes from the respective T cell subsets determined by scRNA-seq.

After scRNA-seq quality control filtering, we performed uniform manifold approximation and projection (UMAP), followed by dimensionality reduction and identified 11 scRNA-seq clusters, which were annotated based on differentially expressed genes (DEGs; log_2_ FC >0.25, Bonferroni adjusted p-value <0.01). In Arm infection, we observed expected T cell phenotypes, including naïve T cells (T^naive^; *Ccr7*, *Sell,* and *Lef1,* 248 DEGs), effector T cells (T^eff^; *Klrg1* and *Ly6c2*, 143 DEGs), effector memory T cells (T^em^; *Klrb1c*, *Klrd1* and *S1pr1*, 193 DEGs), and memory T cells (T^mem^; *Il7r*, *Arl4c* and *Il18r1*, 35 DEGs; **Figure 1F, Table S1**). In Cl13 infection, we observed Tex^prog^ (*Tcf7*, *Slamf6* and *Id3*, 117 DEGs), Tex^int^ (*Lgals3*, *S100a4* and *Mki67*, 113 DEGs), and Tex^term^ (*Gzma*, *Bcl2, Cd101* and *Entpd1*; n=138), as expected (**Figure 1D and 1F**). In addition, we also observed early effector exhausted cells (Tex^eeff^; *Xcl1*, *Top2a* and *Mif*, 1,059 DEGs; a predominant population on D8 of C13 infection), killer cell lectin-like receptor (KLR)-expressing exhausted cells (Tex^KLR^; *S1pr5*, *Cx3cr1*, *Klrc1* and *Zeb2*, 260 DEGs; emerging specifically late in C13), lung terminal exhausted cells primarily detected in the lung (Tex^lung^; *Lag3*, *Ifng*, *Ccl3* and *Ccl4*, 247 DEGs), and interferon signature gene (ISG) exhausted T cells (Tex^ISG^; *Isg15*, *Ifit1*, *Ifit3* and *Isg20*, 273 DEGs; **Figure 1D and F**).

We observed 8 analogous T cell populations in the scATAC-seq data and annotated each cluster based on differential chromatin accessibility at marker gene loci identified in scRNA-seq clusters (i.e., Gene Score, log_2_ FC > 0.5, FDR < 0.01; **Figure 1G**) and integration with scRNA-seq data (**Figure S1F, Methods**). Since our primary goal was to analyze Tex differentiation, we did not perform scATAC-seq at D21 in Arm infection, or in lung or liver T cells in Cl13 infection; thus, scATAC-seq clusters did not include T^em^, T^mem^, or Tex^lung^ subsets. However, scATAC-seq clusters did reveal additional heterogeneity in the early effector subsets, including three early activated/effector populations from the D8 time point in the two infection models. These subsets did not co-cluster with D21 Tex populations, and included two effector populations (T^eff^ and T^eff2^) - mainly derived from the Arm condition - and an early effector exhausted population from the Cl13 condition (Tex^eeff^; **Figure 1E and G, Table S2**).

scATAC-seq profiles were analyzed at the level of: (1) chromatin accessibility of *cis*-elements (open chromatin regions; OCRs), (2) gene activity scores, computed from the accessibility of enhancers linked to a single gene promoter based on proximity and co-accessibility, and (3) transcription factor (TF) activity, computed from the enrichment of TF binding sites in OCRs or the accessibility of TF binding sites genome-wide in each single cell [25, 26]. Analysis of *cis*-elements identified cell type-specific OCRs (T^naive^ - 12,444; T^eff^ - 331; T^eff2^ - 4,070; and Tex^eff^ - 1,463; Tex^prog^ - 4,532; Tex^int^ - 448; Tex^term^ - 2,264; Tex^KLR^ - 668; **Figure 1H, Table S2**), and accessibility was correlated with gene expression at marker gene loci that define Tex subsets, including *Tcf7* (T^naive^, Tex^prog^), *Pdcd1* (Tex^prog^, Tex^int^, and Tex^term^), and *Tox* (Tex^prog^, Tex^int^, and Tex^term^; **Figure 1I**, **Methods**). TF motif enrichment analysis at cell type-specific OCRs identified TF motifs enriched in specific T cell subsets. As expected, T^naive^-specific OCRs were enriched for the TCF/LEF motifs, which were also enriched in Tex^prog^, along with other known Tex^prog^ TFs (e.g., BATF, AP-1 and BACH), and two with undescribed functions (HIVEP and NFKB) [14, 17, 27, 28]. Tex^eeff^ showed NFAT motif enrichment, while KLF motifs were specifically enriched in the Tex^int^, T^eff^, and Tex^KLR^ populations. Finally, Tex^term^-specific OCRs exhibited strong enrichment for NR4A, RUNX and NFAT TF motifs (**Figure S1G, Table S3**) [29–32]. Together, these datasets describe the landscape of transcriptional and epigenetic CD8^+^ T cell states, including previously unidentified Tex populations, that emerge in response to chronic LCMV infection.

### CX3CR1^+^ exhausted T cells comprise three distinct Tex subsets

We first examined heterogeneity within CX3CR1^+^ Tex cells, since these cells have recently been described as a highly proliferative and multi-functional intermediate cell state between Tex^prog^ and Tex^term^ [8, 9, 13]. scRNA-seq of sorted CX3CR1^+^ T cells from D21 of Cl13 infection revealed substantial heterogeneity that primarily spanned three distinct phenotypes (Tex^eeff^, abundant at D8, and Tex^int^ and Tex^KLR^, abundant at D21; **Figure 2A, Figure S2A**). To better understand the temporal gene expression programs of Tex^eeff^ and Tex^int^, we performed DEG analysis (log_2_ FC > 0.25, Bonferroni adjusted p-value < 0.01) and identified 382 genes with significantly higher expression in Tex^eeff^ (e.g., *Rplp0*, *Rpsa*, *Gapdh* and *Cenpa*) and 286 genes with significantly higher expression in Tex^int^ (e.g., *Ccl5*, *Cd3e*, *Lcp2* and *Nfatc1*; **Figure S2A, Table S4**). Ingenuity pathway analysis linked protein translation (EIF2 Signaling), cell cycle (Kinetochore Metaphase Signaling Pathway), and glycolysis (Glycolysis I.) to the D8 population, indicative of a highly proliferative, activated T cell subset. In contrast, the transcriptional program of Tex^int^ at D21 was related to TCR stimulation and downstream signaling (**Figure S2A**). These results suggest that the Tex^eeff^ subset possesses higher cycling and glycolytic activities, while the Tex^int^ subset is more differentiated and expresses genes related to TCR signaling, which seeds downstream Tex populations.

**Figure 2.**
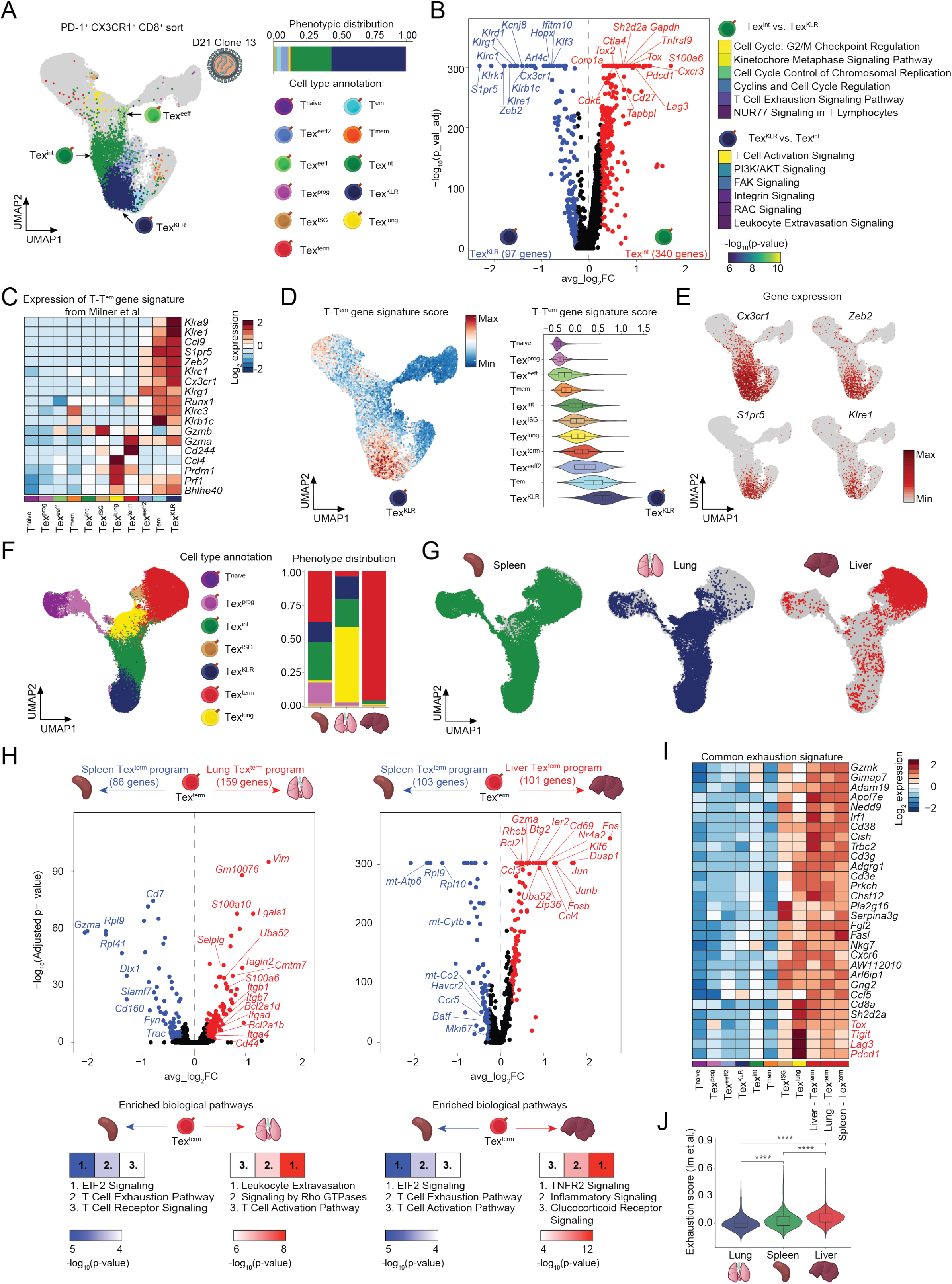
Identification of early effector, KLR-expressing, and organ-specific Tex subsets. **(A)** UMAP of scRNA-seq results colored by the main T cell subsets of the sorted PD-1^+^, CX3CR1^+^ and CD8^+^ T cells. Stacked bar plot shows the phenotypic distribution of the sorted population (right) **(B)** Volcano plot of differentially expressed genes between the Tex^KLR^ and Tex^int^ cell populations (left). Ingenuity pathway analyses on the differentially expressed genes show the enriched biological pathways in the two subsets. Top 6 hits are shown. **(C)** Heatmap of the expression of the marker genes of terminal effector memory (T-T^em^) cells defined by Milner et al. in the indicated T cell subsets. **(D)** UMAP colored by the strength of the T-T^em^ gene signature (T-T^em^ module score) in the scRNA-seq dataset (left). Violin plot representation of the T-T^em^ score in the indicated T cell subsets. **(E)** UMAPs colored by the expression of the indicated marker genes of the T-T^em^ subset. **(F)** UMAP of scRNA-seq results from the three organs at D21 following Cl13 infection colored by the annotated T cell subsets (left). Stacked bar plot representation of the phenotypic distribution of the annotated T cell subsets in the three organs (right). **(G)** UMAPs colored by the cells from the three organs. **(H)** Volcano plots of differentially expressed genes comparing the Tex^term^ cell populations from the different organs. Ingenuity pathway analysis results on the differentially expressed gene groups (bottom). Top 3 hits are shown. **(I)** Heat map of the gene expression values of the common exhaustion gene signature among the organ specific Tex^term^ subsets. **(J)** Violin plot depicts the exhaustion scores of the three organs based on Im et al. 2016.

To similarly determine the transcriptional programs that distinguish Tex^int^ cells from Tex^KLR^ cells, we performed DEG analysis and found 97 Tex^KLR^-biased genes and 340 Tex^int^-biased genes (**Figure 2B, Table S5**). Pathway analysis of Tex^KLR^ and Tex^int^ genes demonstrated the enrichment of cell cycle- and T cell exhaustion-related biological terms in the Tex^int^ population, while linking T cell activation signaling and T cell motility-related functions to the Tex^KLR^ subset **(Figure 2B)**. Notably, many markers of terminal effector and effector memory T cells, such as the killer cell lectin-like receptor (KLR) family members (e.g., *Klrd1*, *Klrk1*, *Klrc1*, *Klre1* and *Klrg1*), the TF, *Zeb2*, and its target gene, *S1pr5* (a marker of tissue emigrating antigen-experienced T cells), showed a highly specific expression pattern in the Tex^KLR^ subset [33, 34]. In contrast, Tex^int^ cells expressed canonical exhaustion markers, such as *Tox*, *Tox2*, *Ctla4*, *Pdcd1*, and *Lag3*, along with cell cycle genes (e.g., *Cdk6*), and TCR signaling genes (e.g., *Coro1a*, *Tapbpl* and *Sh2d2a*; **Figure 2B**) [35–37]. We focused on the Tex^KLR^ subset and assessed the expression of the gene signature of terminal effector memory T cells (T-T^em^), a recently described subset of T^em^ identified during acute LCMV infection, which express effector T cell markers, including KLRs (e.g., *Cx3cr1*, *Zeb2*, *S1pr5*, and *Klre1*) [38]. The T-T^em^ gene signature was highly expressed in the Tex^KLR^ cells and was also observed at the single cell level by scoring the cells based on the expression of this gene panel; thus, cells we term ‘KLR-expressing Tex’ (Tex^KLR^) may represent a parallel differentiation path to T-T^em^ with strong effector function and the potential for memory formation (**Figure 2C-E)**[13]. In summary, the CX3CR1^+^ T cell pool contains additional T cell subsets with distinct functionalities and dynamics during the course of chronic infection, which may explain the multitude of effector- and exhaustion-related functions that have previously been linked to this population [8, 9, 13].

### Tex acquire organ-specific terminal exhaustion signatures

Next, we asked whether chronic viral infection leads to similar T cell states in different tissues. We re-clustered scRNA-seq profiles from animal-matched gp33^+^ and gp33^-^ CD8^+^ T cell fractions from the spleen, lung, and liver at D21 of Cl13 infection. We annotated CD8^+^ T cell subsets based on the previously-defined markers and examined their distribution across organs (**Figure 2F**). Relative to splenic T cells, cells in the lung exhibited an alternative terminal exhausted phenotype (Tex^lung^) and a reduced Tex^prog^ population, while the proportions of Tex^KLR^ and Tex^int^ populations were similar to the spleen (**Figure 2F and G**). Strikingly, T cells in the liver almost exclusively adopted the Tex^term^ phenotype, with dramatically reduced numbers of other Tex phenotypes, as previously described (6.1% of the total**; Figure 2F and G**) [60]. We further examined tissue-specific differences in the exhaustion signature by pairwise differential gene expression analyses (log_2_ FC > 0.25, Bonferroni adjusted p-value < 0.01). Compared to splenic Tex^term^ cells, liver-derived Tex^term^ cells possessed a strong tissue-resident memory T cell signature, including the expression of *Cd69*, *Cxcr6*, *Ccl3* and *Ccl4*, and heightened mTOR and glycolytic activity (**Figure 2H, Table S6**). Similarly, lung-derived Tex^term^ cells also exhibited typical markers of lung-resident memory T cells, including *Cxcr6*, *Cd44* and several integrin genes (*Itga4*, *Itgad*, *Itgab7* and *Itgab1*; **Figure 2H**, **Table S7**). Furthermore, both liver- and lung-derived Tex^term^ cells expressed higher levels of pro-survival genes than splenic Tex^term^ cells, including, *Bcl2*, *Bcl2a1b,* and *Bcl2a1d* (**Figure 2H, Table S8**). These results suggest that Tex^term^ cells can obtain tissue residency signatures and persist in tissues in the setting of chronic antigen.

Despite tissue specific differences in gene expression of Tex^term^, we observed a common Tex^term^ gene signature across all organs. This signature (n=35 genes) contained previously described exhaustion-related genes, such as immune checkpoint inhibitory receptors, *Pdcd1*, *Lag3*, and *Tigit*, and the key TF, *Tox*, which imprints the transcriptional and epigenetic signature of T cell exhaustion (**Figure 2I**). Finally, we constructed an exhaustion gene signature based on previously defined CXCR5^+^ and CXCR5^-^ T cells subsets and scored the severity of exhaustion among Tex^term^ cells from each organ [6]. We observed that liver-derived Tex^term^ cells scored the highest for the exhaustion signature, followed by splenic and lung-derived Tex^term^ cells (**Figure 2J**). We also scored Tex^term^ and Tex^int^ cells based on cell cycle activity, which ranked liver-derived cells as the least proliferative, followed by the lung and spleen, inversely correlating with the severity of exhaustion (**Figure S2B**). These results demonstrate that T cell exhaustion develops across multiple organs with a common gene expression signature but microenvironment-specific effects; namely, exhaustion is most pronounced in the liver niche, which is perhaps driven by higher antigen burden or anatomical differences [39].

### Regulatory programs underlying Tex subsets and early fate commitment to the Tex lineage

The chromatin state of Tex subsets is dynamically regulated and represents a major point of epigenetic imprinting [20] [40]. Two open questions are: (1) the earliest cell cell stage of Tex epigenetic priming, and (2) the temporal regulation of the Tex epigenetic program. To address these questions, we focused our analysis on scATAC-seq data from Cl13 infection and analyzed gp33^+^ and gp33^-^ T cells from two time points (D8 and D21) that encompassed previously-defined and our newly-defined Tex subsets (**Figure 3A and B**). Next, we defined differential OCRs for each Tex subset, including Tex^eeff^ (3,567 OCRs), Tex^prog^ (4,818 OCRs), Tex^int^ (235 OCRs), Tex^KLR^ (1,223 OCRs), and Tex^term^ (1,594 OCRs; **Figure 3C**). Cl13-focused scATAC-seq analysis identified a second, more effector-like Tex^eeff2^ population at D8 that exhibited higher accessibility at *Klrc1* and *Gzmm* genes (2,296 OCRs). In addition to the previously described Tex subset-specific motif enrichments (**Figure S1G**), the Cl13-focused analysis allowed us to observe potential relatedness of the subsets based on their chromatin features and enriched TF motifs **(Figure 3C)**. Namely, we observed that: (1) the open chromatin landscape of Tex^prog^ and Tex^eeff^ partially overlap, indicative of developmental relatedness, and (2) the Tex^int^ subset exhibits an intermediate chromatin state between the Tex^term^ and Tex^KLR^ subsets, with very few unique OCRs, suggesting that this cell state is an intermediate cell stage and a potential bifurcation point of Tex differentiation, supported by our observation of Tex^int^ at D8 and the emergence of Tex^term^ and Tex^KLR^ by D21 (**Figure 1B**).

**Figure 3.**
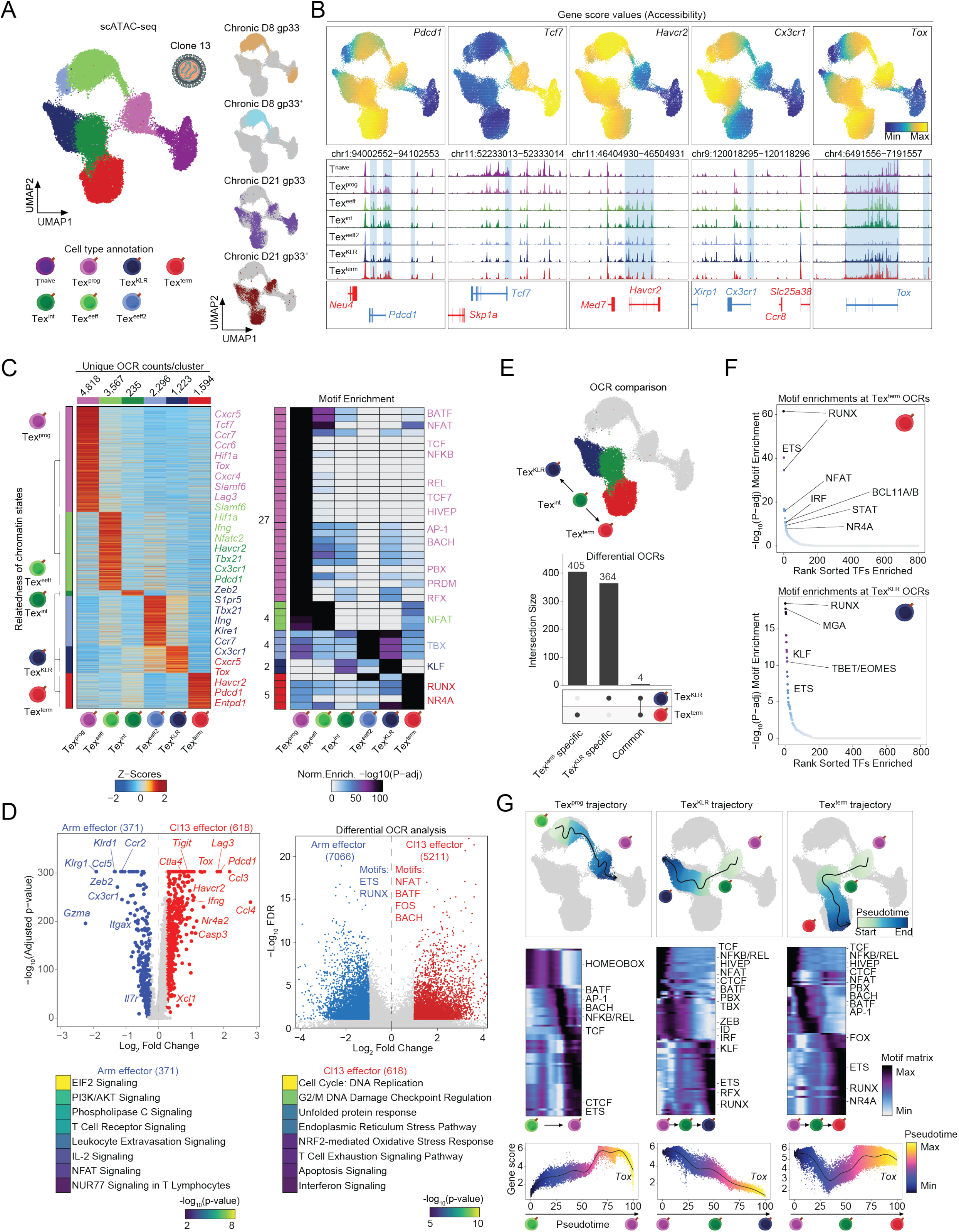
Tex^int^ represent a bifurcation point of exhausted T cell fate differentiation. **(A)** UMAP of scATAC-seq results of D8 and D21 gp33^+^ and gp33^−^ T cells from the Cl13 infection model. UMAP is colored by the annotated T cell subsets. Small UMAPs (right) show T cells that originate from the indicated gp33 fractions and timepoints. **(B)** Feature plots of the indicated Gene score values (accessibility) (top) and genome browser snapshots of the corresponding genomic loci (bottom). **(C)** Heat map of Peak score values at the unique open chromatin regions (OCRs) of the T cell subsets determined by scATAC-seq with a list of annotated putative target genes based on proximity (left). Heat map of motif enrichment results at the unique OCR sets of the annotated T cell subsets. **(D)** Volcano plot of differentially expressed genes between the Arm effector cells and Cl13 early effector cells (left). Ingenuity pathway analysis results show the top 8 enriched biological terms (bottom). Volcano plot depicts the differentially accessible OCRs between the Arm effector cells and Cl13 early effector cells (right). **(E)** UMAP depicts the populations used for differential OCR analysis (top). Upset plot of differentially accessible OCRs and their overlap among the Tex^KLR^ and Tex^term^ populations (bottom). **(F)** Hockey stick plots depict the enriched transcription factor motifs at the specific OCRs of the Tex^term^ and Tex^KLR^ subsets. **(G)** Pseudotime trajectory analyses of three potential Tex differentiation paths (top). Heat maps show transcription factor deviation scores that change over the pseudotime trajectories (middle). Gene score values of *Tox* on the three pseudotime trajectories (bottom).

Next, we focused on the early programming of exhaustion by comparing D8 scATAC-seq phenotypes in Arm and Cl13 infection **(Figure S3A)**. As previously described, memory precursor cells (T^mp^) are present at D8 in Arm infection and cluster with an early Tex^prog^ population present at D8 in Cl13 infection that expresses *Tox* and *Tcf7* (herein referred to as precursor exhausted - Tex^prec^), but these subsets were relatively infrequent compared to the effector populations in both infection models (1.4% of D8 cells in Arm infection, 3.3% of cells at D8 in Cl13 infection) [14, 41]. We first compared the gene expression and chromatin state programs of T^mp^ and Tex^prec^ subsets, which revealed strong exhaustion- and interferon-induced programs in Tex^prec^, as expected (**Figure S3B, S3C and S3D, Table S9**). Second, we analyzed DEGs of effector cells in Arm and Cl13 infection, which revealed a strong Tex signature in the Tex^eeff^ subset compared to T^eff^; T^eff^ showed a *bona fide* effector program (e.g., *Gzma*, *Klrd1*, *Ccr2*, and *Cx3cr1*, 371 DEGs), while Tex^eeff^ expressed high levels of exhaustion marker genes (e.g., *Tox*, *Lag3*, *Pdcd1*, *Havcr2*, *Ctla4*, and *Tigit,* 618 DEGs; **Figure 3D**). These observations were also supported by the chromatin state programs of these subsets (T^eff^ - 7,066 OCRs vs. Tex^eeff^ - 5,211 OCRs) that were associated with T^eff^-specific (ETS and RUNX) and Tex^eeff^-specific (NFAT and BATF) TF motifs (**Figure 3D, Table S10**). Altogether, these results support recent studies demonstrating the formation of Tex^prec^ early during chronic infection that exhibit molecular signatures of exhaustion, distinct from T^mp^ [14, 41]. However, we find that the exhaustion program, including *Tox* expression, is present in an earlier Tex^eeff^ stage and driven by NFAT and BATF, which may prime the chromatin state of TCF1^-^ cells for exhaustion, supporting a model in which an initial wave of effector cells undergoes contraction and gives rise to Tex^prog^ cells that seed additional Tex subsets.

Although Tex differentiation downstream of Tex^prog^ is currently thought to follow a linear path, our identification of a Tex^KLR^ subset, which emerges late in infection alongside Tex^term^, suggests that the Tex^int^ population may represent a potential bifurcation point between Tex^KLR^ and Tex^term^ phenotypes (**Figure 1D, 2A and 3A**) [3]. We compared scATAC-seq profiles of Tex^KLR^ and Tex^term^ subsets to Tex^int^ cells and identified 405 Tex^term^-specific, 364 Tex^KLR^-specific OCRs, and only 4 common OCRs, suggesting that these two cell states are epigenetically divergent (**Figure 3E**). Accordingly, TF motif enrichment analysis showed increased accessibility of NFAT, IRF, STAT, and NR4A TF motifs in Tex^term^ and increased accessibility of RUNX, MGA, KLF, TBET/EOMES and ETS TF motifs in Tex^KLR^ (**Figure 3F**). Differential gene expression analysis (log_2_ FC > 0.25, Bonferroni adjusted p-value < 0.01) identified 97 Tex^KLR^-biased genes (e.g., *Klrg1*, *Arl4c* and *Zeb2*) and 340 Tex^term^-biased genes (e.g., *Tox*, *Tox2*, *Lag3* and *Pdcd1*; **Figure S3E, Table S11**). These results indicate that Tex^KLR^ and Tex^term^ cells exhibit distinct chromatin and gene expression programs, supporting the idea that these phenotypes represent late stages of a divergent differentiation trajectory of exhaustion that bifurcates at the Tex^int^ stage.

Finally, we analyzed 15,809 variable OCRs for TF motif enrichments across three differentiation trajectories nominated by longitudinal timepoint data and/or chromatin state similarities: (1) Tex^prog^ trajectory (Tex^eeff^ → Tex^prec^ → Tex^prog^), (2) Tex^term^ trajectory (Tex^prog^ → Tex^int^ → Tex^term^), and (3) Tex^KLR^ trajectory (Tex^prog^ → Tex^int^ → Tex^KLR^; **Figure 3G**). The Tex^prog^ trajectory showed a gradual loss of HOMEOBOX TF motifs and enrichment of BATF, AP-1, BACH, NFKB, TCF and CTCF motifs. In contrast, in both Tex^term^ and Tex^KLR^ trajectories, we observed a gradual loss of Tex^prog^ specific TF motifs (e.g., TCF, BACH and BATF) upon entry to the Tex^int^ cell state. Differentiation trajectories that bifurcated from the Tex^int^ state showed the enrichment of specific TF motifs that might bind TFs which can guide the differentiation program of Tex^KLR^ (e.g., ZEB, ID, IRF, KLF, ETS, RFX, HIVEP and RUNX) and Tex^term^ (e.g., RUNX and NR4A; **Figure 3G**). Finally, we studied the accessibility of the *Tox* locus, encoding the TF that is critical for Tex differentiation [4, 15, 16, 18, 19]. Accessibility of the *Tox* locus gradually increased as cells transitioned from Tex^eeff^ to Tex^prog^, while it gradually decreased as they transitioned to Tex^KLR^. The Tex^term^ trajectory demonstrated a decrease in *Tox* accessibility during the Tex^prog^ to Tex^int^ transition and a subsequent increase in the Tex^term^ state (**Figure 3G**). We annotated differentially accessible OCRs (compared to T^naive^ cells) in a +/− 250kb window around the transcription start site of *Tox* and identified 88 OCRs. Of these OCRs, 16 and 8 were differentially accessible in Tex^eeff^ or Tex^prog^, respectively, which was also supported by high *Tox* expression in these subsets (relative to T^naive^), indicating that TOX executes the molecular programming of Tex differentiation in these subsets (**Figure S3F, Table S12**). These results identify the Tex^eeff^ population as a novel point of the molecular programming of exhaustion and nominate the Tex^int^ population as a potential bifurcation point of Tex cell differentiation states.

### Clone tracing reveals divergent Tex differentiation trajectories during chronic viral infection

We next leveraged paired scRNA/TCR-seq data to analyze clonal trajectories of T cells in Arm and Cl13 (D8 and D21) infection (**Figure 4A**). We identified 212 and 280 expanded T cell clones (> 1 cell) at D8 and D21 of Arm infection, respectively, and 134 and 338 expanded clones at D8 and D21 of Cl13 infection, respectively. As expected, at D8 of Arm infection, clonally expanded T cells were largely restricted to the T^eff^ pool, while at D21, clonally expanded T cells exhibited a balanced distribution between T^em^ and T^mem^ phenotypes (**Figure 4B and Figure S4A**). In contrast, clonal expansion at D8 in Cl13 infection occurred almost exclusively in Tex^eeff^ **(Figure 4C)**. Importantly, the Tex^prog^ population did not show strong clonal expansion at this early time point, further supporting our prior observation that cells in this population are infrequent at D8 and subsequently expand by transition from TEX-eeff or self-renewal from cells not significantly present at D8 **(Figure 3)**. At D21 in Cl13 infection, we observed expanded clones across multiple Tex phenotypes, including Tex^prog^, Tex^int^, Tex^term^, and Tex^KLR^ (**Figure 4C and Figure S4B**).

**Figure 4.**
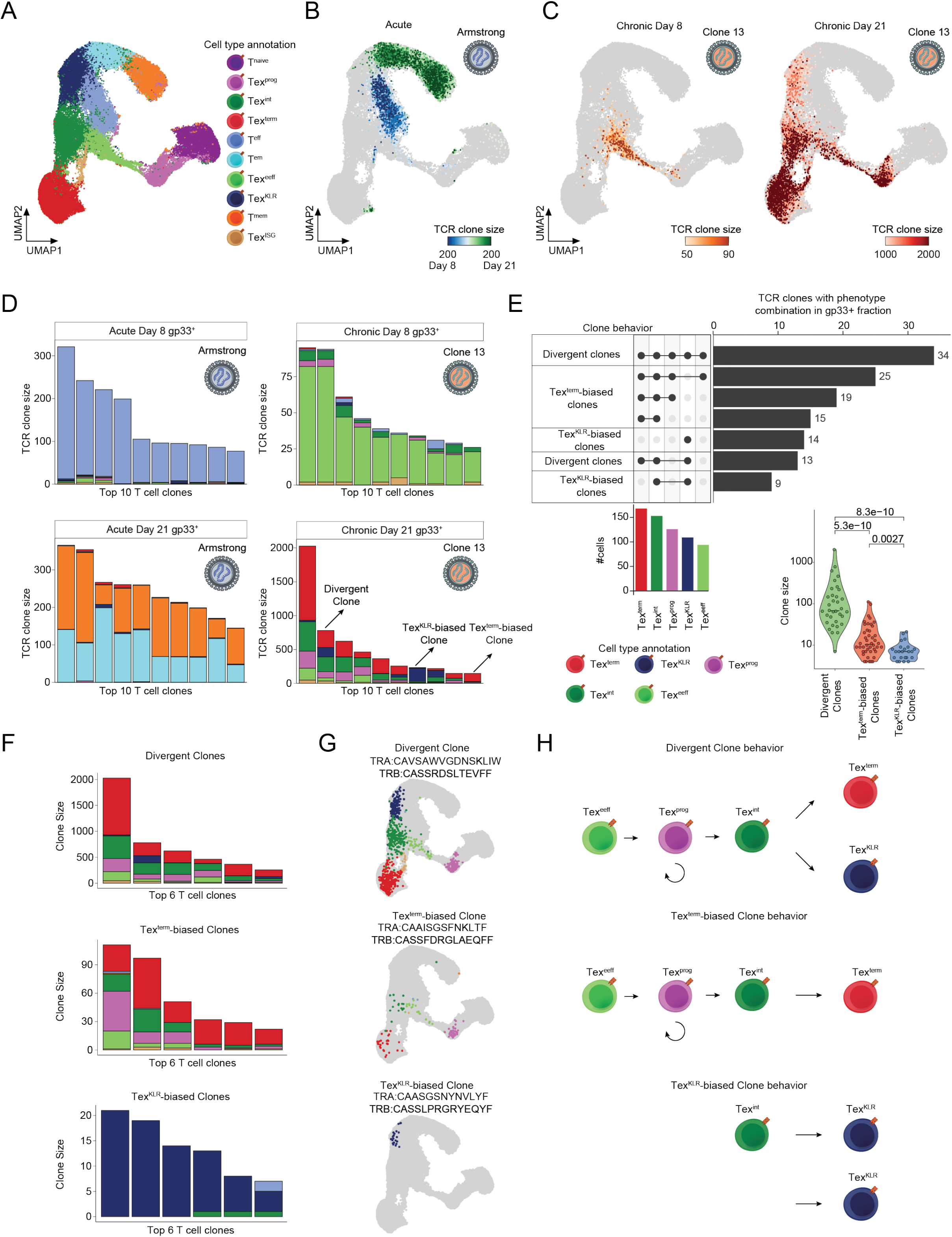
TCR-based lineage tracing reveals divergent Tex clonal trajectories. **(A)** UMAP of scRNA-seq results from the gp33^+^ and gp33^−^ T cell factions of the Arm and Cl13 infection model from D8 and D21 following infection. UMAP is colored by the annotated T cell subsets. **(B)** UMAP of scRNA-seq results colored by the size of the detected TCR clones at D8 and D21 in the Arm infection model. **(C)** UMAP of scRNA-seq results colored by the size of the detected TCR clones at D8 in the Cl13 infection model (left). Same UMAP colored by the TCR clone size at the D21 time point in the Cl13 infection model (right). **(D)** Stacked bar plot of the phenotypic distribution of the top 10 expanded clones in the gp33^+^ fraction of Arm D8 and D21 samples (left). Same stacked bar plots representing the top 10 expanded clones in the Cl13 infection model (right). **(E)** Upset plot depicting the expanded clones with specific phenotype combinations (clone behaviors). Barplot shows the number of cells with the indicated phenotypes that make up the expanded clones. Violin plot shows the clone size distribution of the detected clone behaviors. **(F)** Stacked bar plots show the top 6 expanded clones with the indicated clone behaviors. **(G)** UMAPs show representative examples for the detected clone behaviors. **(H)** Scheme on the phenotypic composition and the potential differentiation trajectories of the identified clone behaviors.

To further investigate Tex clonal differentiation trajectories, we visualized the distribution of cellular phenotypes for the top 10 expanded clones at D8 and D21 in each infection. At D8, cells of the top expanded clones from the Arm condition almost exclusively acquired the T^eff^ phenotype, with clone sizes ranging from 77-321 cells (mean 153 cells, 3.8% of 4,030 total cells). At D21, top expanded clones acquired both T^em^ and T^mem^ phenotypes, with clone sizes ranging from 145-366 (mean 246 cells, 3.5% of 7,033 total cells; **Figure 4D and Figure S4A**). In contrast, the top expanded clones in Cl13 infection acquired the Tex^eeff^ phenotype at D8, with clone sizes ranging from 26-95 cells (mean 49 cells, 3.5% of 1,414 total cells). Notably, expanded clones only contained small numbers of cells with the Tex^prog^ phenotype at this time point (**Figure 4D**). Analysis of D21 of Cl13 infection identified substantially larger clone sizes, ranging from 146-2,026 cells (mean 525 cells, 7.0% of 7,489 total cells; **Figure 4D**). Strikingly, these large clones contained cells with multiple Tex phenotypes (Tex^prog^, Tex^int^, Tex^KLR^, and Tex^term^), although the frequency of each phenotype varied considerably between individual clones. Namely, individual clones either preferentially acquired the Tex^term^ or the Tex^KLR^ phenotypes, or developed into both phenotypes (**Figure 4D and Figure S4B**). This observation prompted us to perform a more detailed analysis of the phenotypic distribution of all large clones (> 3 cells detected) of the top 7 most dominant clonal phenotype combinations, which revealed three main clonal differentiation patterns (referred to as clone behaviors): 1) Tex^term^-biased clones, consisting of cells that predominantly acquired the Tex^term^ and not Tex^KLR^ phenotype (45% of clones), 2) Tex^KLR^-biased clones, consisting of T cells that predominantly acquired the Tex^KLR^ phenotype (18% of clones), and 3) divergent clones, consisting of cells that acquired Tex^term^ and Tex^KLR^ phenotypes (37% of clones; **Figure 4E**). Divergent clones were the most clonally expanded and ranged from 7-2,026 cells (mean 197 cells) per clone, while Tex^term^-biased clones ranged from 4-111 cells (mean 19 cells) per clone. Interestingly, Tex^KLR^-biased clones were relatively small and ranged from 4-21 cells (mean 8 cells) per clone (**Figure 4E-G**). We also noted several larger clones (4-233 cells, mean 53 cells) that skewed heavily to the Tex^KLR^ phenotype (>50% of cells acquire the Tex^KLR^ phenotype), but had a small percentage of Tex^term^ cells (**Figure S4C**). To account for sampling biases where not all relevant phenotypes may be observed for small clones, we randomized T cell phenotype and TCR clone assignment to generate a null distribution of clone patterns if each clone randomly acquired all observed phenotypes (**Methods**). This analysis revealed a striking enrichment of Tex^KLR^- and Tex^term^-biased clone behavior over random chance, whereas the divergent clonal differentiation pattern was twice as likely to be detected by random chance than observed in our data, suggesting that the observed biases in clone behavior are not simply the result of sampling bias (**Figure S4D**). Altogether, these results reveal novel clonal Tex differentiation trajectories during chronic infection **(Figure 4H**).

### Antigen-specific expanded Tex clones and phenotypes are shared across tissues

Next, we asked if clonal differentiation patterns are intrinsically programmed, perhaps by the TCR, or stochastic. We first determined whether expanded Tex clones could be found across different tissues by analyzing antigen-specific gp33^+^ and gp33^−^ CD8^+^ T cells across organs (animal-matched) in Cl13 at D21 (**Figure 5A**). In spleen-, liver-, and lung-derived scRNA/TCR-seq datasets, we detected expanded T cell clones across all three tissues, and as expected, the gp33^+^ and gp33^−^ fractions showed minimal TCR overlap, validating our sorting strategy (**Figure 5B and Figure S5A**). Importantly, there was significant TCR sharing across the different organs for both gp33^+^ and gp33^−^ fractions (**Figure S5A**). We identified expanded organ-shared T cell clones that had at least 5 T cells, which consisted of at least 1 cell from each organ. This analysis identified 100 shared T cell clones among all organs, 37 clones shared between the lung and spleen, and 22 clones specific to the spleen (**Figure 5C and D**). We examined the degree of expansion of TCR clones that were detected across organs and observed a strong correlation in clone frequency in each pairwise organ comparison (spleen:liver - *R*=0.66; spleen:lung - *R*=0.65; liver:lung - *R*=0.76, **Figure 5C**).

**Figure 5.**
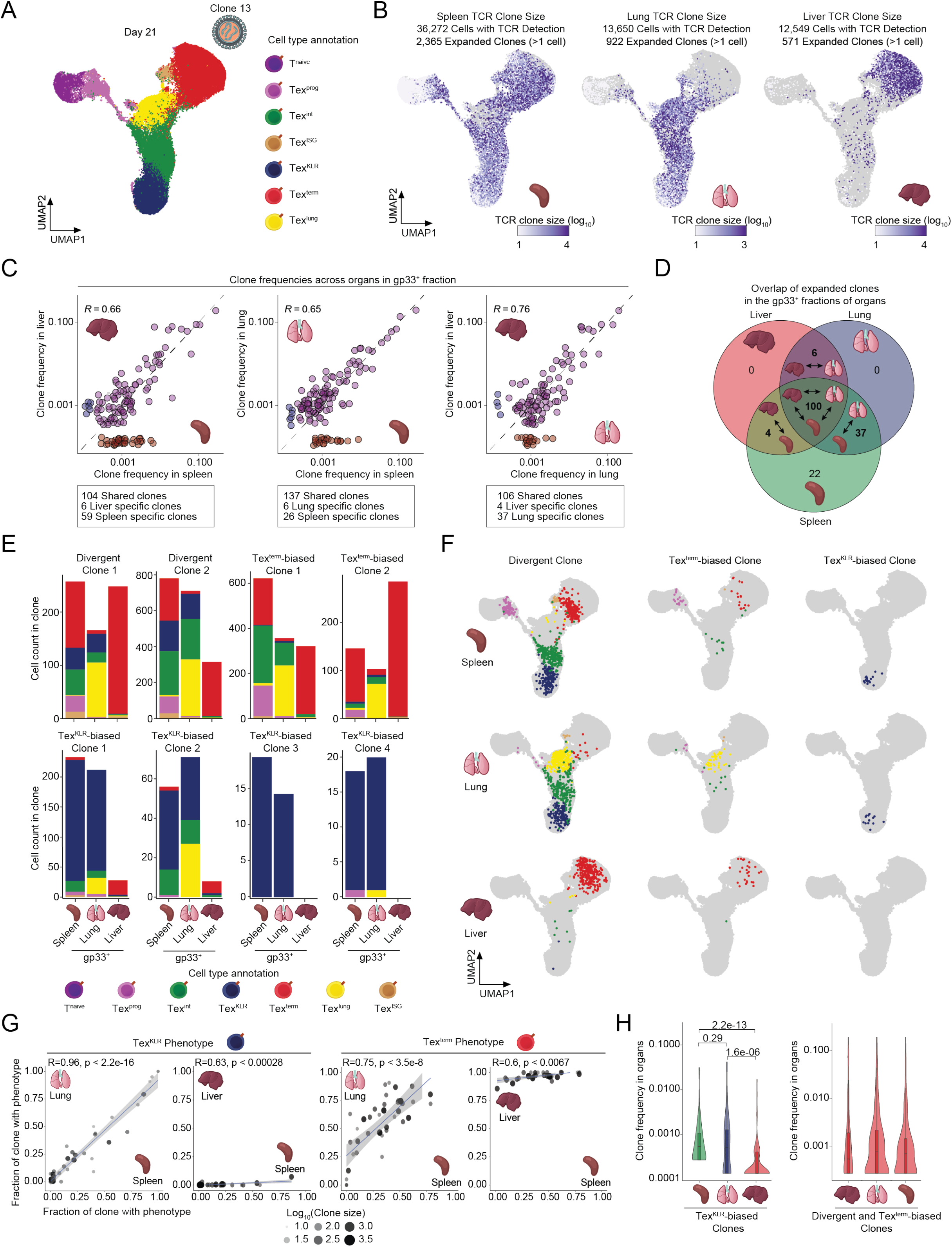
Conserved clonal T cell trajectories across organs and depletion of Tex^KLR^ in the liver microenvironment. **(A)** UMAP of organ-derived T cells at D21in Cl13 infection colored by the annotated T cell subsets. **(B)** UMAPs colored by the detected TCR clone sizes in the different organs. **(C)** Scatterplots depicting the frequencies of expanded T cell clones from the indicated organ comparisons. The correlation coefficient, and specific and shared clone numbers are indicated for each comparison. **(D)** Venn diagram depicting the overlap of expanded T cell clones in the gp33^+^ fraction of the indicated organs. **(E)** Stacked bar plot of the phenotypic composition of individual clones across organs. **(F)** UMAPs depict individual clones with specific clone behaviors among organs. **(G)** Scatter plots showing the fraction of the shared clones with Tex^KLR^ and Tex^term^ phenotypes between the indicated organs. **(H)** Violin plot depicts Tex^KLR^-biased clone frequencies across the organs, which includes clones with >50% Tex^KLR^ phenotype (left). Violin plot of Tex^term^-biased and divergent clone frequencies across the organs.

Next, we examined the distribution of phenotypes for clones shared across organs with different differentiation trajectories defined by their trajectory in the spleen (**Figure 5E, F and Figure S5B, C**). First, we focused on comparisons between the spleen and lung, since they exhibited similar heterogeneity in Tex phenotypes. Strikingly, differentiation trajectories were highly conserved between the two tissues. Divergent clones in the spleen also maintained Tex^term^ and Tex^KLR^ phenotypes in the lung (although instead exhibiting the aforementioned terminal Tex^lung^ phenotype; 35/48 divergent clones detected in both organs, **Figure 5E, F, Figure S5B, C**). Similarly, the majority of splenic Tex^KLR^-biased clones remained Tex^KLR^-biased in the lung (4/7 clones detected in both organs) and the majority of splenic Tex^term^-biased clones remained Tex^term^-biased in the lung (11/14 clones detected in both organs; **Figure 5E and Figure S5C**). In particular, we did not observe appreciable interconversion between Tex^KLR^- and Tex^term^-biased clones between these two organs (0/19 shared clones). Accordingly, quantification of the Tex^KLR^ and Tex^term^ frequencies within individual clones showed a high concordance across organs (Tex^KLR^ spleen:lung - *R*=0.96, Tex^term^ spleen:lung - *R*=0.75; **Figure 5G**). Altogether, these results demonstrate that clonally expanded Tex clones are shared across organs and that clonal differentiation behavior is primarily an intrinsically programmed, rather than stochastic, process.

### Depletion of Tex^KLR^ clones in the liver microenvironment

We next analyzed clonal behavior in the liver, which showed an enrichment of Tex^term^ compared to other organs (94% Tex^term^), perhaps driven by high antigen burden [39]. Thus, in contrast to the lung, we expected an enrichment in clonal Tex^term^ frequency; however, this could either be driven by: (1) depletion of Tex^KLR^ in the liver microenvironment, or (2) interconversion of Tex^KLR^-biased clones to Tex^term^-biased clones. To distinguish between these two possibilities, we first analyzed the Tex^KLR^-biased clones from the spleen and found that only one of these clones was present in the liver (1/7 shared clones), suggesting that Tex^KLR^-biased clones are depleted in the liver niche. Similarly, although divergent clones were largely detectable in the liver (52/58 clones shared between spleen and liver), we again observed a depletion of Tex^KLR^ cells, resulting in Tex^term^-biased behavior in the majority of the cases (32/52 shared clones). In contrast, the majority of Tex^term^-biased clones remained Tex^term^-biased in the liver, although they were heavily skewed towards the Tex^term^ phenotype, with relative loss of Tex^prog^ and Tex^int^ phenotypes (9/9 clones, **Figure 5E, F and Figure S5C**). Importantly, Tex^term^-biased clones did not adopt a Tex^KLR^ phenotype. Quantification of frequencies of Tex^KLR^ and Tex^term^ phenotypes of shared clones in the spleen and liver confirmed the depletion of Tex^KLR^ cells in the liver and a skewing of Tex^term^-biased clones to the Tex^term^ fate **(Figure 5G and H)**. Altogether, these results demonstrate that Tex clones entering the liver exhibit changes in clonal behavior due to the loss of Tex^KLR^, suggesting that Tex^KLR^ are not able to persist in high antigen environments, perhaps due to activation-induced cell death.

### TCR affinity can program Tex clone behavior and phenotypic fate commitment

The difference in expansion levels between Tex^term^-biased clones and Tex^KLR^-biased clones led us to examine whether Tex differentiation trajectories were driven by differences in TCR affinity. We used tetramer staining as a proxy for TCR affinity against the immunodominant LCMV epitope, gp33, and sorted gp33^−^ (n=8,914), gp33-intermediate (gp33^int^; n=5,875), and gp33-high (gp33^high^; n=8,194) CD8^+^ T cells from the spleen of Cl13-infected mice at D21 and performed scRNA/TCR-seq (**Figure 6A and Figure S6A**). Analysis of TCR sequences identified 313 TCRs in gp33^high^ cells, 1,576 TCRs in gp33^int^ cells, and 3,803 TCRs in gp33^−^ cells (**Figure S6B**). The TCR repertoire showed a relatively small overlap between gp33^high^ and gp33^−^ cells (13 shared TCRs), compared to the overlap between gp33^high^ and gp33^int^ cells (158 shared TCRs), or gp33^int^ and gp33^−^ cells (306 shared TCRs), and quantification of TCR repertoire similarity using the Morisita overlap index demonstrated that gp33^int^ sorting captured a distinct TCR repertoire compared to gp33^high^ and gp33^−^ fractions (**Figure 6B and Figure S6B**).

**Figure 6.**
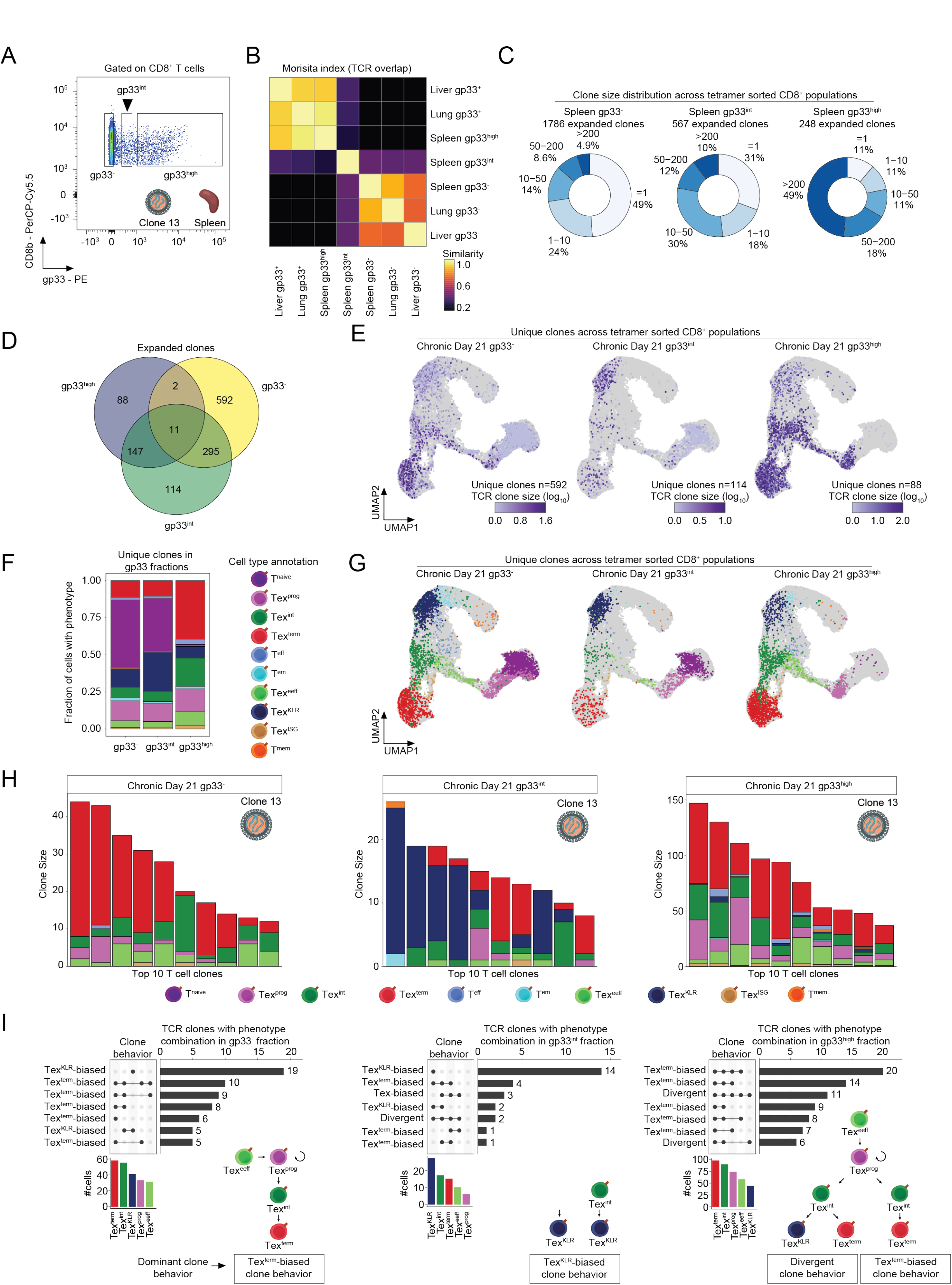
Differences in TCR signal strength regulate clonal differentiation of Tex^KLR^ and Tex^term^. **(A)** Sorting strategy to obtain gp33^−^, gp33^int^ and gp33^high^ T cell populations from the spleen of LCMV-Cl13 infected animals 21 days following infection. **(B)** Heat map depicting TCR repertoire overlap (Morisita index) among the different gp33 fractions from the indicated samples. **(C)** Pie chart representation of the fraction of the detected clone sizes in the three gp33 T cell fractions. **(D)** Venn diagram depicts the overlap of the expanded clones from the gp33 T cell fractions. **(E)** UMAPs colored by size of the unique expanded clones in the three gp33 T cell fractions. **(F)** Stacked bar plot of the phenotypic distribution of the unique expanded clones of the three gp33 T cell fractions. **(G)** UMAPs visualizing the unique expanded clones of the three gp33 T cell fractions colored by the annotated T cell subsets. **(H)** Stacked bar plot of the top 10 uniquely expanded T cell clones from the three gp33 T cell fractions colored by the annotated T cell phenotypes. **(I)** Upset plots depict the unique expanded clones with specific phenotype combinations (clonotype behavior) from the three gp33 T cell fractions. Barplots show the number of cells with the indicated phenotypes. Dominant clone behaviors are indicated at the bottom.

Next, we evaluated the clone size distribution of the sorted populations, which revealed an increase in the percentage of large clones (clones with 5-200 or >200 cells) as a function of higher tetramer fluorescence, with an accompanying decrease in clonal diversity (**Figure 6C**). To link unique TCRs to each gp33-tetramer fraction, we compared the overlap of clones between gp33 fractions and identified 592 unique gp33^−^ clones, 114 unique gp33^int^ clones, and 88 unique gp33^high^ clones (**Figure 6D**). Importantly, the distribution of cellular phenotypes for these unique clones showed considerable phenotypic skewing (**Figure 6E-G**). Namely, gp33^high^ cells contained ~3.3 times more cells with Tex^term^ and Tex^int^ phenotypes, compared to either gp33^−^ or gp33^int^ cells (39% Tex^term^ and 19% Tex^int^ in gp33^high^; 11% Tex^term^ and 6.7% Tex^int^ in gp33^int^; 11% Tex^term^ and 7.0% Tex^int^ in gp33^−^), indicating a pronounced phenotypic skewing towards terminal exhaustion. In contrast, gp33^int^ cells exhibited phenotypic skewing towards the Tex^KLR^ phenotype in the population, compared to the gp33^high^ and gp33^−^ populations (27% Tex^KLR^ in gp33^int^; 7.9% Tex^KLR^ in gp33^high^; 13% Tex^KLR^ in gp33^−^; **Figure 6F and G**).

To further analyze differentiation trajectories at a clonal level, we visualized the top 10 unique expanded clones in each gp33-tetramer fraction and assessed their phenotypic composition. We found that the top clones in the gp33^−^ and gp33^high^ fractions were biased towards Tex^term^ or divergent phenotypes (10/10 gp33^−^ clones and 10/10 gp33^high^ clones), while in contrast, the largest clones in the gp33^int^ pool exhibited phenotypic skewing towards the Tex^KLR^ phenotype (5/10 gp33^int^ clones; **Figure 6H**). Finally, we analyzed the clone behaviors of the unique clones of the three gp33 fractions in the top 7 most dominant phenotypic patterns that define clone behaviors (**Figure 6I**). Clones of the gp33^−^ fraction exhibited two major clone behaviors, Tex^term^-biased and Tex^KLR^-biased. Interestingly, clones from the gp33^int^ fraction were heavily enriched for Tex^KLR^-biased differentiation. Finally, expanded clones unique to the gp33^high^ fraction were biased towards Tex^term^-biased and divergent clone behaviors, and no Tex^KLR^-biased clones were identified **(Figure 6I**). Surprisingly, divergent clone behaviors were much more common in the unique T cell clones of the gp33^int^ and gp33^high^ fractions compared to gp33^−^ clones, suggesting that this differentiation path may be more common among T cell clones that recognize this immunodominant epitope. These results establish that T cell clones distinguished by their affinity for the immunodominant LCMV epitope have divergent differentiation paths, with lower affinity TCR clones favoring the development of Tex^KLR^ and higher affinity TCR clones biasing toward Tex^term^ and divergent behavior.

## Discussion

Here we report a single-cell multi-omic atlas of T cell exhaustion during chronic viral infection, which reveals novel Tex subsets, identifies multiple differentiation trajectories of Tex clones, and nominates TCR signal strength as a key driver of clonal behavior. We define an early effector Tex differentiation state (Tex^eeff^), where the molecular program of exhaustion is initiated, and identify a bifurcation point of Tex differentiation (Tex^int^), which can give rise to two alternative late-stage Tex phenotypes (Tex^KLR^ and Tex^term^) with the potential to balance effector function, immunological memory, and persistence in high antigen environments. Using the TCR sequence as an endogenous molecular barcode, we track the fate of individual T cell clones and establish three main clonal developmental trajectories that give rise to the heterogeneous Tex pool. Surprisingly, we find that clonal differentiation patterns are shaped by TCR affinity and affect the resulting phenotype and clonal expansion in different tissue microenvironments. These findings highlight the importance of studying the polyclonal T cell repertoire at single cell resolution to fully uncover the diversity and function of T cell states in the immune response.

Prior studies have described multiple Tex subsets with distinct phenotypic and functional traits, primarily within the spleen microenvironment during chronic viral infection [3,10, 11, 42, 43]. In addition to Tex^term^ and Tex^prog^ subsets, transitory exhausted cells have more recently been characterized as a multi-functional CX3CR1^+^ population with high cytolytic activity, proliferative capacity, and the ability to contribute to the memory T cell pool [8, 9, 13]. Here we show that this CX3CR1^+^ population encompasses three T cell subsets with distinct functionalities: 1) an early effector exhausted subset (Tex^eeff^) with high proliferative capacity early in infection that is largely absent at later stages; 2) intermediate exhausted T cells (Tex^int^), which maintain a high proliferation signature and upregulate signaling downstream of TCR stimulation, and 3) a Tex^KLR^ subset with a strong cytolytic gene expression program, and a terminal effector memory cell-like signature that has been described in acute infection [38].

Given the distinct, stable epigenetic state of Tex, which persists after antigen clearance [20, 40, 44–46], a key question is the stage at which Tex epigenetic imprinting occurs. Previous studies have shown that early TCF1^+^ Tex^prec^ cells possess the epigenetic signature of Tex and can seed additional Tex subsets [14, 41]. Here, we find that the Tex program is initiated at an earlier stage in TCF1^−^ Tex^eeff^. scATAC-seq analysis suggests that this fate decision is initially driven by NFAT and BATF, which may prime the chromatin state of TCF1^−^ cells to develop into Tex^prec^, which subsequently activate BACH2 and TCF-1 to give rise to Tex^prog^ [14, 17]. This finding supports a model in which the Tex^prec^ pool, and eventually the Tex^prog^ pool originates from Tex^eeff^, analogous to memory differentiation from memory precursors or short-lived effector cells during acute infection.

Downstream of the Tex^prog^ population, the differentiation trajectory of Tex has largely been shown to follow a linear cellular path [3]. However, our data suggests that there are two late-stage cell types that result from a divergent differentiation path (Tex^KLR^ and Tex^term^), and that individual clones can follow three differentiation trajectories resulting in Tex^term^-biased, Tex^KLR^-biased, or divergent fates, comprising both cell types. Furthermore, we find that the differentiation trajectory of Tex clones is intrinsically programmed by TCR affinity and conserved across specific tissue microenvironments; high-affinity TCR clones are biased towards divergent and Tex^term^ differentiation trajectories, while low-affinity TCR clones are biased towards a Tex^KLR^ trajectory. However, the presence of clones with divergent behavior suggests that there may be additional paths to induce TCR signal strength variation – perhaps via inhibitory receptor signaling, access to antigen, antigen-presenting cell type, or other factors – to generate Tex^KLR^. Importantly, Tex^KLR^-biased clones were dramatically depleted in the liver microenvironment, suggesting that these clones and this phenotype are sensitive to the antigen-rich environment of the liver and are unable to persist. Given the high viral load and inflammatory microenvironment of the liver during infections, these results suggest that the Tex^term^ phenotype precludes activation-induced cell death, improves Tex persistence, and preserves anti-viral effector function in the organ system [47].

Finally, these findings may have several implications for cancer, where T cell exhaustion can limit the T cell response and efficacy of immunotherapies. First, several ongoing therapeutic strategies aim to reverse exhaustion; however, our results suggest that Tex^term^ may be specifically adapted to survive in high antigen niches, and that inhibiting Tex^term^ differentiation may be deleterious, rather than beneficial, to the T cell response [4, 15, 16, 18, 19, 48]. Whether the pro-survival aspects of T cell exhaustion can be specifically maintained, while still reinvigorating other aspects of effector function will require further study. Second, our findings reinforce the notion that TCR signal strength directs the phenotypic fate of T cells, in addition to mediating recognition of specific antigens [49, 50]. Thus, the generation of TCR-based cellular therapies should incorporate the assessment of phenotypic outcomes of TCR activation, in addition to peptide-MHC binding properties. Finally, the observation that a polyclonal T cell response to chronic antigens balances persistence, effector, and potential memory functions via the development of two Tex states suggests that future cellular therapies should also aim to establish divergent phenotypes, encompassing Tex^term^ and Tex^KLR^. Future studies should investigate whether Tex^KLR^ develop during tumor-specific T cell responses. A recent study identified a natural killer (NK) cell-like signature in chronic antigen-induced exhausted human chimeric antigen receptor (CAR)-T cells, which resembles the Tex^KLR^ signature described here, suggesting that this cell type may be present in adoptive cell therapy settings as well [51]. Manipulation of these Tex states and their underlying gene regulatory programs and differentiation pathways may provide avenues to improve T cell-based immunotherapies in the future.

## Acknowledgements

We thank the members of the Satpathy, Egawa and Chang labs for stimulating discussions. This work was supported by the National Institutes of Health (NIH) K08CA230188 (A.T.S.), U01CA260852 (A.T.S.), RM1-HG007735 (H.Y.C.), R01AI130152 (T.E.), R21AI161040 (T.E.), a Career Award for Medical Scientists from the Burroughs Wellcome Fund (A.T.S.), a Technology Impact Award from the Cancer Research Institute (A.T.S.), an ASH Scholar Award from the American Society of Hematology (A.T.S.), the Parker Institute for Cancer Immunotherapy (H.Y.C., and A.T.S.), a Leukemia and Lymphoma Society Scholar Award (T.E.), and the Scleroderma Research Foundation (H.Y.C.). H.Y.C. is an investigator of the Howard Hughes Medical Institute. K.E.Y. was supported by the National Science Foundation Graduate Research Fellowship Program (NSF DGE-1656518), a Stanford Graduate Fellowship and a NCI Predoctoral to Postdoctoral Fellow Transition Award (NIH F99CA253729). J.A.B was supported by a Stanford Graduate Fellowship and a National Science Foundation Graduate Research Fellowship under Grant No. DGE-1656518. The sequencing data was generated with instrumentation purchased with NIH funds: S10OD018220 and 1S10OD021763.

## Author contributions

B.D., K.E.Y. and A.T.S. conceptualized the study. B.D., K.E.Y. and A.T.S. wrote and edited the manuscript and all authors reviewed and provided comments on the manuscript. B.D., K.S., K.E.Y., K.J.H.G., X.Y. and Y.Q. performed experiments. K.E.Y., S.L.M. and J.A.B. analyzed data. B.D., K.E.Y., J.R.G., E.J.W., H.Y.C., T.E. and A.T.S. guided experiments and data analysis.

## Data availability

Reviewer access for sequencing data is available under GEO accession: GSE188670.

## Methods

### Mice and infection

Male C57BL/6N mice were purchased from Charles River Laboratories. All mice were housed in a specific pathogen-free facility at Washington University in St. Louis and were used for infection at 8–12 week of age. LCMV infection was performed essentially as described previously [52]. All experiments were performed according to a protocol approved by Washington University’s Institutional Animal Care and Use Committee.

### Tissue preparation

Single cell suspension of the different organs was prepared by manual dissociation. Organs were minced and gently pushed through a 40-micron strainer. Spleen single cell suspensions were spun, and red blood cells were lysed with ACK-lysis buffer by resuspending the cell pellet followed by 2 minutes incubation. Cells were then washed with ice-cold PBS and stained for sorting in FACS buffer (PBS, 0.1% BSA, 2mM EDTA, 5% FBS). For the lung and liver single-cell suspension, organs were cut into small pieces and gently pushed through a 40-micron diameter strainer. Single-cell suspensions were then layered on top of Ficoll-Paque Plus (Cytiva) and centrifuged according to the manufacturer’s recommendations. The lymphocyte fraction was collected and washed with ice-cold PBS, and then stained for sorting.

### Staining T cells for sorting

Single cell suspensions were stained with the following antibodies: CD8b (PerCP-Cy5.5), PD-1 (PE-Cy7), CX3CR1 (APC), SLAMF6 (BV605) and the class I tetramer, H-2Db LCMV gp33-41 (KAVYNFATC) (PE). Cells were stained with the tetramer for 20 minutes at 4C followed by staining with the combination of the other antibodies for 20 minutes. Cells were washed in FACS buffer and stained with LIVE/DEAD Fixable Aqua dead cell stain for 20 minutes in PBS.

### scATAC-seq sample and library generation

Single cell ATAC-seq experiments were performed on the 10x Chromium platform as described earlier [53]. Briefly, after sorting, T cells were washed with PBS + 0.04% BSA and then subjected to nuclei isolation according to the protocol of the manufacturer. Nuclei were counted and on average ~10,000 nuclei were submitted for tagmentation. After tagmentation, nuclei were loaded for capture using the 10x Chromium controller. After Gel emulsion generation, linear amplification was performed, followed by DNA purification according to the manufacturer’s protocol. The resulting DNA was used for library construction as described on the website of the manufacturer. Libraries were quantified by Agilent Bioanalyzer and were sequenced on an Illumina NovaSeq S4 sequencer, using the following setup: 50bp read 1N, 8bp i7 index, 16bp i5 index and 50bp read 2N. In this reaction, 1N and 2N refers to the DNA insert sequencing, while i5 and i7 sequencing identifies the individual barcodes of single cells.

### Single-cell RNA-seq library preparation

Single-cell RNA-seq libraries were prepared using the 10X 5’ Single Cell Immune Profiling Solution Kit (v1.1 Chemistry), according to the manufacturer’s instructions. Briefly, FACS sorted cells were washed once with PBS + 0.04% BSA and on average 10,000 cells were submitted for capture using the 10x Chromium controller. Following reverse transcription and cell barcoding in droplets, emulsions were broken, and cDNA was purified using Dynabeads MyOne SILANE followed by PCR amplification (98°C for 45 sec; 14 cycles of 98°C for 20 sec, 67°C for 30 sec, 72°C for 1 min; 72°C for 1 min). For gene expression library construction, 50 ng of amplified cDNA was fragmented, end-repaired, and double-sided size selected with SPRIselect beads. Purified DNA was subjected to PCR amplification with sample indexing primers (98°C for 45 sec; 14 cycles of 98°C for 20 sec, 54°C for 30 sec, 72°C for 20 sec; 72°C for 1 min). Amplified DNA was double-sided size selected with SPRIselect beads and were quantified using Agilent Bioanalyzer. Single-cell RNA-seq libraries were sequenced on an Illumina NovaSeq S4 sequencer using the following read configuration 26bp Read1, 8bp i7 Index, 91bp Read2.

### Single-cell TCR library generation

Single-cell TCR libraries were prepared with the 10x Chromium Single Cell V(D)J Enrichment Kit for mouse T cells (v1.1 Chemistry) following the manufacturer’s protocol. Briefly, after cDNA amplification and clean up, 2ul of cDNA was used for target enrichment. First, target enrichment 1 was performed by specific primers followed by a SPRIselect bead clean-up. Second, target enrichment 2 was performed with specific primers followed by double-sided size selection with SPRIselect beads. After the two target enrichment steps, the quality of the product was assessed with Agilent Bioanalyzer. Amplified product was then subjected for fragmentation, followed by end repair and A-tailing. End repaired product was then subjected to adaptor ligation followed by SPRIselect bead purification. Product was amplified and barcoded with adaptor specific primers and the quality of the resulting libraries were determined by Agilent Bioanalyzer. Single-cell TCR-seq libraries were sequenced on an Illumina NovaSeq S4 sequencer using the following read configuration 26bp Read1, 8bp i7 Index, 91bp Read2.

### scATAC-seq data processing and analysis

scATAC-seq datasets were processed as described previously [54]. Briefly, reads were filtered, trimmed, and aligned to the mm10 reference genome using 10X Genomics’ cellranger-atac count pipeline (version 1.2.0).

Processed fragment files were loaded into ArchR (version 1.0.1) for additional processing and analysis. All functions used default parameters unless otherwise specified. Cells were filtered during Arrow file generation using ArchR’s createArrowFiles function to remove cells with an enrichment of Tn5 insertions in transcription start sites (TSS enrichment) of less than 4 or less than 1000 unique fragments. Doublets were identified using ArchR’s addDoubletScores function and predicted doublets removed using ArchR’s filterDoublets function. Dimensionality reduction was performed using Iterative Latent Semantic Indexing (LSI) using ArchR’s addIterativeLSI function. After initial clustering and UMAP projection, we excluded a small cluster of non-T cells. Cell clustering was performed using ArchR’s addClusters function on IterativeLSI reduced dimensions 1:10 and a resolution of 0.4 (reducedDims = “IterativeLSI”, dimsToUse = 1:10, resolution = 0.4). The same dimensions were used for single cell embedding by Uniform Manifold Approximation and Projection (UMAP) using ArchR’s addUMAP function using IterativeLSI reduced dimensions 1:10 and a minimum distance of 0.1 (reducedDims = “IterativeLSI”, dimsToUse = 1:10, minDist = 0.1). Cell clustering and UMAP projection for Chronic LCMV (D8 and D14, Figure 3) and Day 8 (Chronic and Acute, Figure S3B) subsets were performed as described above with the following modifications: dimsToUse = NULL, resolution = 0.2, and minDist = 0.4.

GeneScore matrices were computed by summing Tn5 insertions in the gene promoter and gene body during Arrow file generation using ArchR’s createArrowFiles function [54]. Gene score imputation was performed with Magic using ArchR’s addImputeWeights function [55]. After clustering the cells, peaks were called by MACS2 on pseudoreplicates sampled from each cluster to obtain a reproducible peak set retaining cell type specific peaks using ArchR’s addReproduciblePeakSet function. Peak co-accessibility and Peak2Gene linkages were computed using ArchR’s addCoAccessibility and addPeak2GeneLinks functions. Transcription factor (TF) motif; deviations were computed with chromVar using ArchR’s addDeviationsMatrix function [26]. Pseudo-bulk tracks for indicated groups of cells were plotted using ArchR’s plotBrowserTrack function with default normalization method based on reads in transcription start sites (“ReadsInTSS”). Differential peak testing was performed using ArchR’s getMarkerFeatures function with testMethod = “wilcoxon” and bias = c(“TSSEnrichment”, “log10(nFrags)”). TF motif enrichment in differential peavks was performed using ArchR’s peakAnnoEnrichment function. Trajectory analysis was performed using ArchR’s addTrajectory and plotTrajectory functions. Identification of positive TF regulators was performed using ArchR’s correlateMatrices function to examine the correlation between chromVar deviation z-scores of TF motifs (“MotifMatrix”) and imputed gene expression (“GeneIntegrationMatrix”) following cross-platform linkage with scRNA-seq data using ArchR’s addGeneIntegrationMatrix.

### scRNA-, TCR-seq computational methods

scRNA-seq reads were aligned to the mm10 reference genome and quantified using cellranger count (10x Genomics, version 3.1.0). Filtered gene-barcode matrices that contained only barcodes with unique molecular identifier (UMI) counts that passed the threshold for cell detection were used for further analysis. scTCR reads were aligned to the mm10 reference genome and consensus TCR annotation was performed using cellranger vdj (10x Genomics, version 3.1.0). TCR annotation was performed using the 10x cellranger vdj pipeline as described.

Additional analysis was performed in R (version 4.0.3) using Seurat (version 4.0.1) using default function parameters unless otherwise noted [56]. Doublets were predicted using DoubletFinder (version 2.0.3) [57]. Cell types were predicted using SingleR (version 1.4.1) based on mouse bulk RNA-seq reference data (MouseRNAseqData) from celldex (version 1.0.0) [58]. Cells with less than 200 genes detected, greater than 5% mitochondrial RNA content, predicted doublets from DoubletFinder, and cells annotated as non-T and non-NK cells by SingleR were excluded from analysis. We predicted cell cycle phase based on previously defined gene sets using the CellCycleScoring function [59]. We then split cells by experimental batch and cell cycle (non-cycling or G1 vs. cycling or G2M/S) into four datasets using Seurat’s SplitObject and performed batch correction using Seurat’s reciprocal PCA workflow. First, we normalized and identified variable features for each dataset independently using Seurat’s NormalizeData and FindVariableFeatures. Then we selected variable features across datasets using Seurat’s SelectIntegrationFeatures. We excluded variable TCR (^Tr.v) genes, variable Ig (^Ig.v) genes, cell cycle genes (used for cell cycle scoring), and mitochondrial genes (^mt-) from integration features used for downstream analysis. We then scaled data and ran PCA on each dataset independently using these features using Seurat’s ScaleData and RunPCA. We identified integration anchors using Seurat’s FindIntegrationAnchors using non-cycling datasets as reference datasets and rpca for dimensionality reduction. We integrated all datasets using Seurat’s IntegrateData using dims=1:50. Integrated data was used for data scaling with ScaleData and PCA dimensionality reduction with RunPCA. After initial clustering we noted three small clusters representing 7% of total cells which had low number of genes detected and high mitochondrial RNA content which were excluded from further analysis. Clusters were identified using shared nearest neighbor (SNN) based clustering based on the first 15 PCs with resolution = 0.45. The same principal components were used to generate the UMAP projections, which were generated with a minimum distance of 0.1. Cell clustering and UMAP projection for Chronic Day 21 T cells (all tissues, Figure 2 and Figure 5), spleen derived T cells (Chronic and Acute, Day 8 and Day 21, Figure 4 and Figure 6), and Day 8 T cells (Spleen, Chronic and Acute, Supplemental Figure 3) were performed as described above with the following modifications:

> Chronic Day 21 T cells: dims = 1:10, resolution = 0.25, min.dist = 0.1
>
> Spleen derived T cells: dims = 1:8, k.param = 50, resolution = 0.45, min.dist = 0.1
>
> Day 8 T cells: dims = 1:12, k.param = 40, resolution = 0.28, min.dist = 0.2

Expression of selected genes was plotted using log normalized gene expression values based on original RNA count data prior to data integration. Marker genes were identified using Seurat’s FindAllMarkers using a cutoff of p_val_adj < 0.01. Differential gene expression analysis was performed using Seurat’s FindMarkers using a cutoff of p_val_adj < 0.05 and abs(avg_log2FC) > 0.25. Gene module scoring was performed using Seurat’s AddModuleScore. TCR clone behaviors were visualized using UpSetR (version 1.4.0). Null distribution of TCR clone behaviors was determined by randomly shuffling TCR clonotype and scRNA phenotype and generating a distribution of TCR clone phenotype combinations (n=50 iterations). Morisita-Horn index for quantifying TCR overlap was calculated using the mh function from the R package divo (version 1.0.1).

---

**Figure S1.**
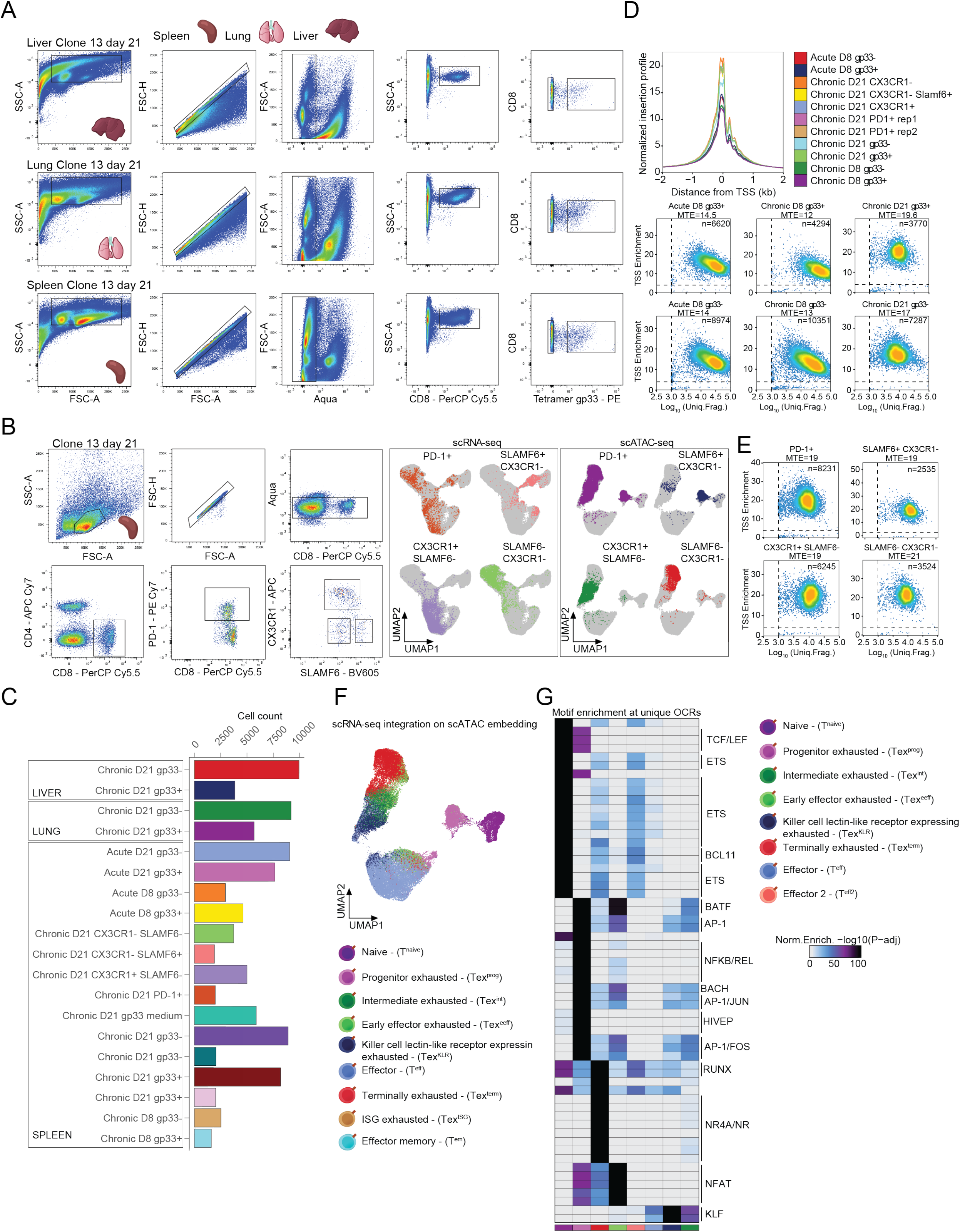
Sorting strategy and quality controls of scATAC-seq data. Related to Figure 1. **(A)** Sorting strategy to obtain antigen specific gp33^+^ and gp33^-^CD8^+^ T cells from different organs. **(B)** Sorting strategy to obtain the main exhausted T cell subsets (left). UMAPs of scRNA-seq and scATAC-seq results, originating from the main, indicated exhausted T cell subsets. **(C)** Bar plot representation of cell counts from the scRNA-seq results. **(D)** Quality control of scATAC-seq data. Histogram shows normalized read enrichment on the transcription start sites (TSS) of genes from the indicated samples (top). Density plots depict the cells that passed the TSS enrichment and Log_10_ unique fragment count threshold. Median TSS enrichment (MTE) is also indicated. **(E)** Density plots of scATAC-seq data from the main exhausted T cell populations depicting the same quality controls as on panel C. **(F)** UMAP of scATAC-seq data colored by the integrated scRNA-seq cluster labels. **(G)** Heat map of motif enrichments at the specific open chromatin regions (OCRs) of the annotated T cell populations.

**Figure S2.**
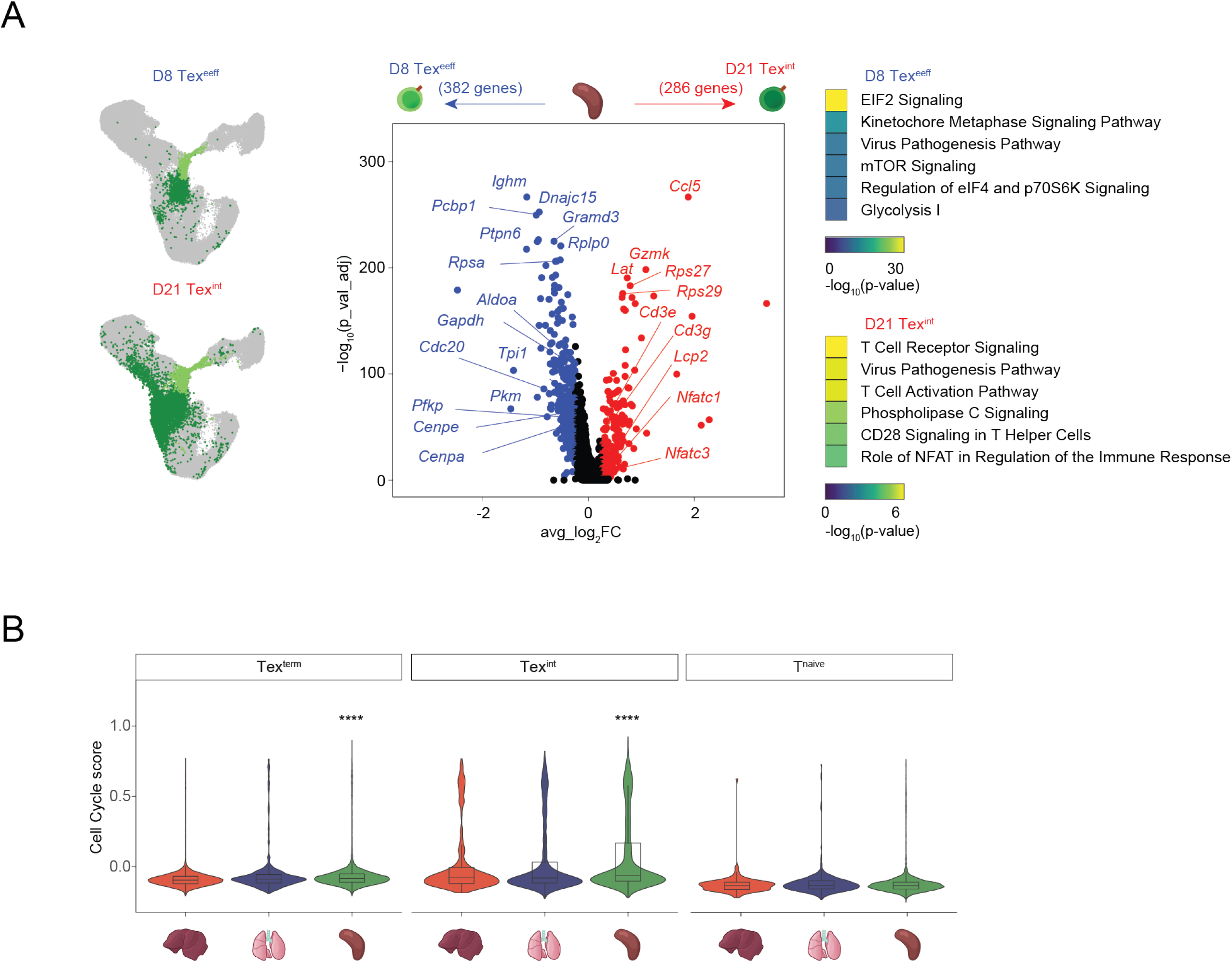
Early effector exhausted cells and intermediate exhausted T cells are phenotypically different populations with distinct temporal appearance. Related to Figure 2. **(A)** UMAPs visualize the early effector exhausted population at D8 and the intermediate exhausted population at D21 (left). Volcano plot depicts the differentially expressed genes between the two populations (middle). Ingenuity pathway analysis results depict the top 6 enriched biological terms in the two populations. **(B)** Violin plots depict the Cell Cycle score of the indicated T cell populations across the indicated organs.

**Figure S3.**
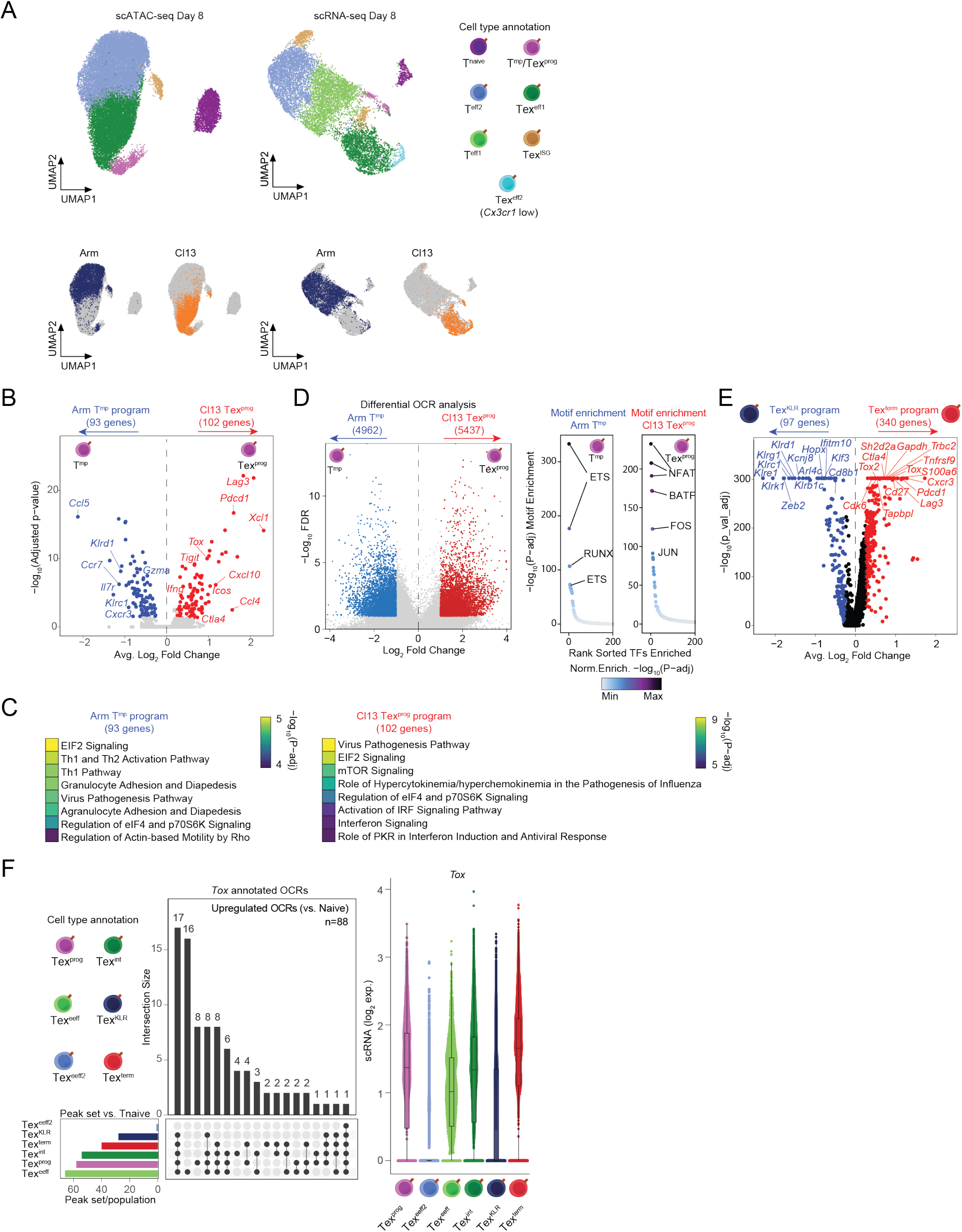
Early progenitor exhausted T cells possess the molecular program of exhaustion. Related to Figure 3. **(A)** UMAPs depict scATAC-seq (left) and scRNA-seq (right) results from the D8 Arm and Cl13 infections. Cells on the small UMAPs are colored by their origin from the two infection models (bottom). **(B)** Volcano plot of differentially expressed genes between the memory precursor T cells (T^mp^) of the Arm and the progenitor exhausted T cells (Tex^prog^) of the Cl13 infection model. **(C)** Ingenuity pathway analyses of the T^mp^ and Tex^prog^ specific gene sets. Top 8 enriched biological terms are shown. **(D)** Volcano plot depicts the differential open chromatin regions (OCRs) of the T^mp^ and Tex^prog^ populations (left). Hockey stick plots show the enriched transcription factor motifs at the specific OCR sets of the T^mp^ and Tex^prog^ subsets. **(E)** Volcano plot of the differentially expressed genes between the Tex^KLR^ and Tex^term^ subsets. **(F)** Upset plot of differentially accessible OCRs annotated to the *Tox* gene relative to T^naive^ cells and their overlap among the different Tex cell subsets. Violin plot shows the gene expression level of *Tox* in the identified Tex subsets.

**Figure S4.**
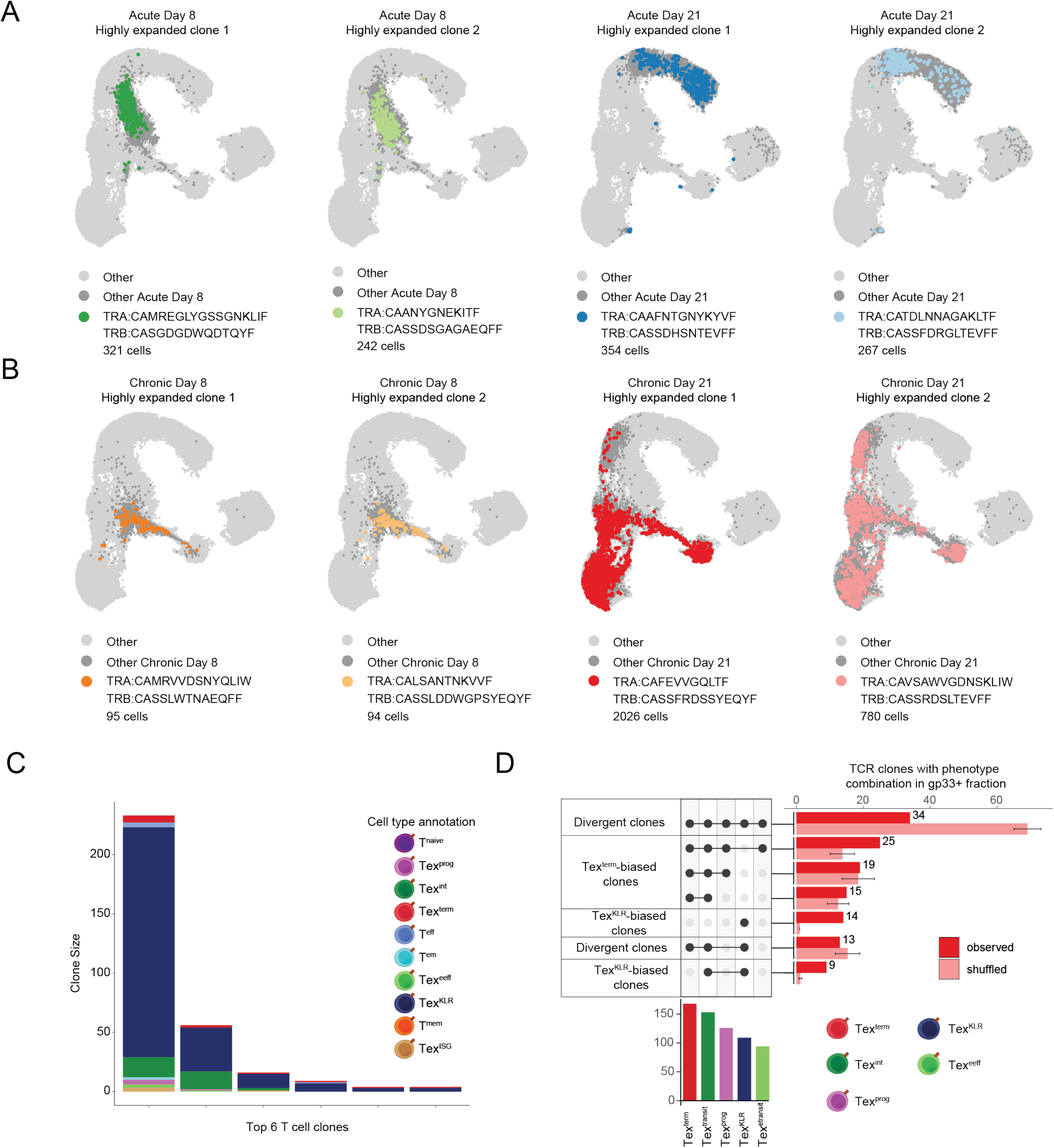
Highly expanded clones of the Arm and Cl13 infection model describe the dominant clone behaviors of exhausted T cell differentiation. Related to Figure 4. **(A)** UMAPs depict highly expanded clones from the Arm infection model at the indicated time points. **(B)** UMAPs depict highly expanded clones of the Cl13 infection model at the indicated time points. **(C)** Stacked bar plot of the phenotypic composition of individual T cell clones with a bias towards the Tex^KLR^ fate, but also exhibiting the Tex^term^ phenotype. Top 6 clones are shown. **(D)** Upset plot of the phenotype combinations of the observed and shuffled TCR clones.

**Figure S5.**
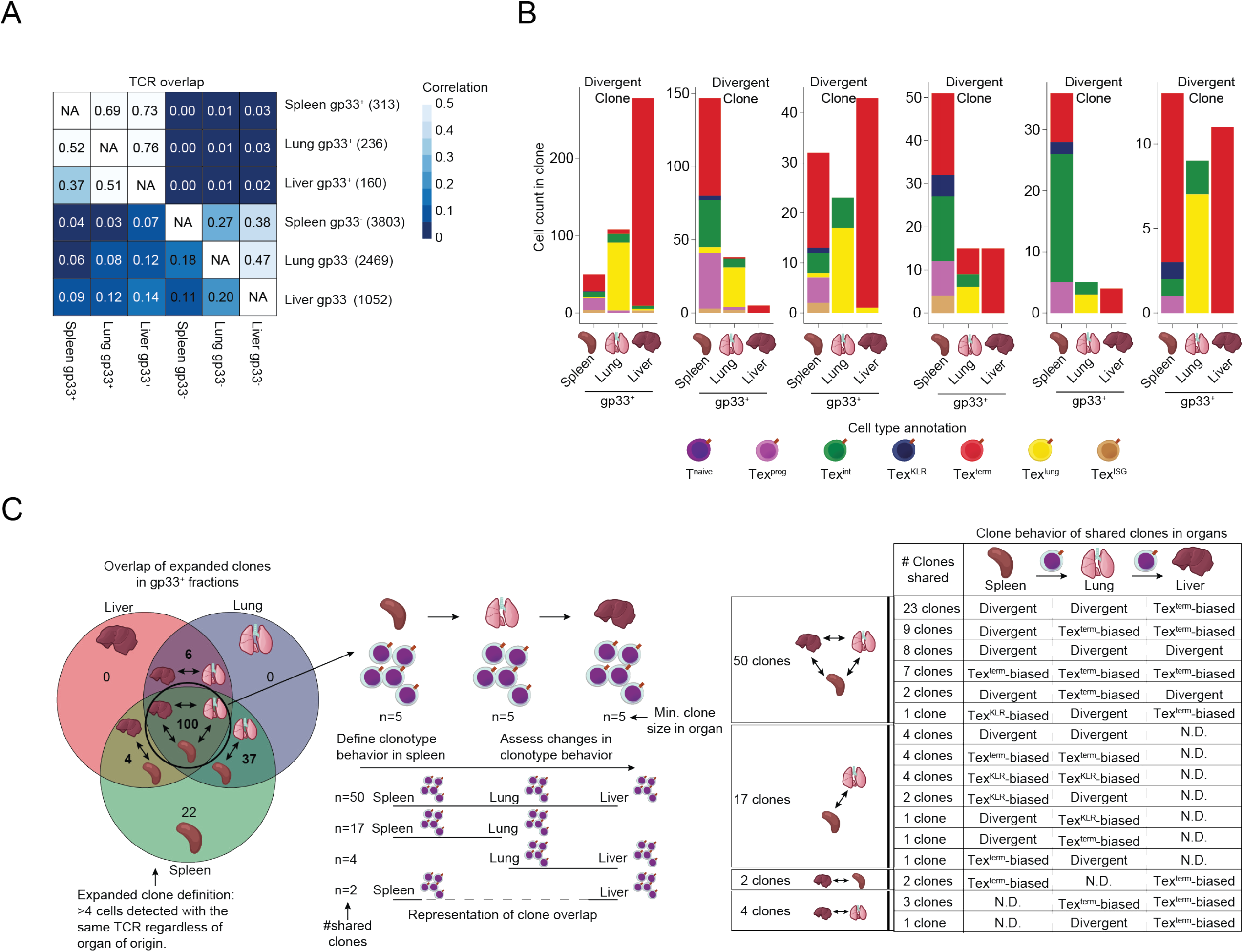
scRNA/TCR-seq reveals the clone behaviors of different organs. Related to Figure 5. **(A)** Heat map representation of the correlation between the TCR repertoires of the indicated gp33^+^ and gp33^-^CD8^+^ T cell subsets from different organs. **(B)** Stacked bar plot of the phenotypic composition of individual clones across organs. **(C)** Schematics show the definition of an expanded, organ-shared T cell clone for clone behavior analysis. Only those clones were considered that had at least 5 T cells present in each organ. Shared clone numbers across the organs are indicated (left). Table depicting the number of expanded clones that are shared across tissues and their clone behaviors (right).

**Figure S6.**
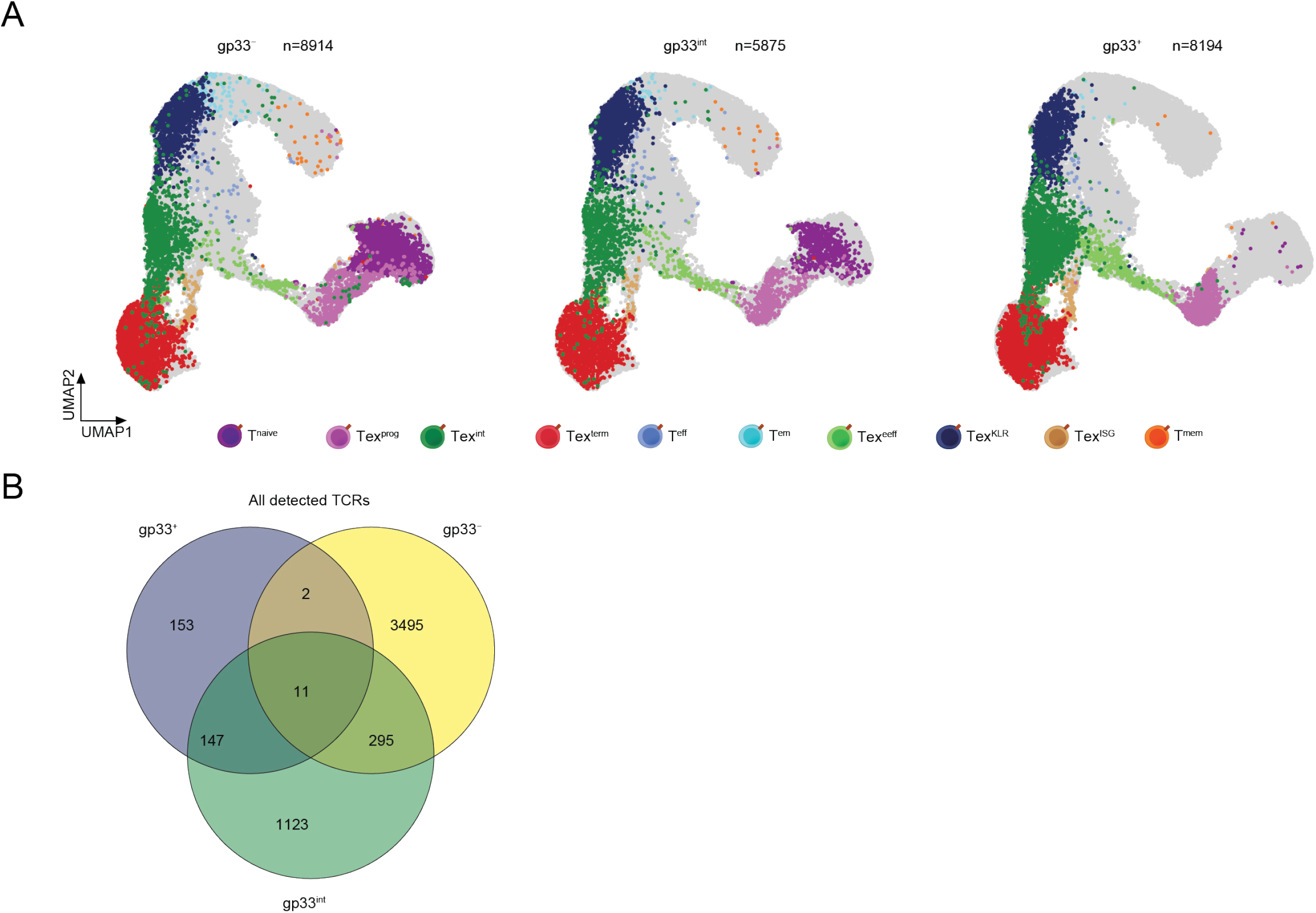
scRNA-seq reveals the phenotypic composition of T cell subsets with different affinities to recognize the immunodominant viral epitope. Related to Figure 6. **(A)** UMAPs of scRNA-seq results colored by the phenotypic distribution of the three gp33 fractions of T cells. **(B)** Venn diagram shows the overlap of all detected TCR clones among the three gp33 T cell fractions.

